# Newfound coding potential of transcripts unveils missing members of human protein communities

**DOI:** 10.1101/2020.12.02.406710

**Authors:** Sebastien Leblanc, Marie A Brunet, Jean-François Jacques, Amina M Lekehal, Andréa Duclos, Alexia Tremblay, Alexis Bruggeman-Gascon, Sondos Samandi, Mylène Brunelle, Alan A Cohen, Michelle S Scott, Xavier Roucou

**Affiliations:** Department of Biochemistry and Functional Genomics, Université de Sherbrooke, Sherbrooke, Quebec, Canada; PROTEO, Quebec Network for Research on Protein Function, Structure, and Engineering; Department of Family Medicine, Université de Sherbrooke, Sherbrooke, Quebec, Canada

**Keywords:** alternative proteins, protein network, protein-protein interactions, pseudogenes, affinity purification-mass spectrometry

## Abstract

Recent proteogenomic approaches have led to the discovery that regions of the transcriptome previously annotated as non-coding regions (i.e. UTRs, open reading frames overlapping annotated coding sequences in a different reading frame, and non-coding RNAs) frequently encode proteins (termed alternative proteins). This suggests that previously identified protein-protein interaction networks are partially incomplete since alternative proteins are not present in conventional protein databases. Here we used the proteogenomic resource OpenProt and a combined spectrum- and peptide-centric analysis for the re-analysis of a high throughput human network proteomics dataset thereby revealing the presence of 280 alternative proteins in the network. We found 19 genes encoding both an annotated (reference) and an alternative protein interacting with each other. Of the 136 alternative proteins encoded by pseudogenes, 38 are direct interactors of reference proteins encoded by their respective parental gene. Finally, we experimentally validate several interactions involving alternative proteins. These data improve the blueprints of the human protein-protein interaction network and suggest functional roles for hundreds of alternative proteins.

## Introduction

Cellular functions depend on myriads of protein-protein interactions networks acting in consort, and understanding cellular mechanisms on a large scale will require a relatively exhaustive catalog of protein-protein interactions. Hence, there have been major efforts to perform high throughput experimental mapping of physical interactions between human proteins (Luck *et al*, 2017). The methodologies involve binary interaction mapping using yeast 2-hybrid (Rolland *et al*, 2014), biochemical fractionation of soluble complexes combined with mass spectrometry (Wan *et al*, 2015), and affinity-purification coupled with mass-spectrometry (Huttlin *et al*, 2015, 2017; Liu *et al*, 2018).

In parallel to these experimental initiatives, computational tools were developed to help complete the human interactome (Keskin *et al*, 2016). Such tools are particularly useful for the identification of transient, cell-type or environmentally dependent interactions that escape current typical experimental protocols. Computational methods that can be used at large scales are created and/or validated using protein-protein interactions previously obtained experimentally (Keskin *et al*, 2016; Kovács *et al*, 2019). Thus, although computational tools complement experimental approaches, the experimental detection of protein-protein interactions is key to building a comprehensive catalog of interactomes.

The BioPlex network is the largest human proteome-scale interactome; initially, BioPlex 1.0 reporting 23744 interactions among 7668 proteins was followed by BioPlex 2.0, which forms the basis of the current study, with 56553 interactions reported involving 10961 proteins. Recent pre-print BioPlex 3.0 reached 118162 interactions among 14586 proteins in HEK293T cells (Huttlin *et al*, 2017, 2015, 2020). The enrichment of interactors of roughly half of currently annotated (or reference) human proteins allowed the authors to functionally contextualize poorly characterized proteins, identify communities of tight interconnectivity, and find associations between disease phenotypes and these protein groups. Here, a community represents a group of nodes in the network that are more closely associated with themselves than with any other nodes in the network as identified with an unsupervised clustering algorithm. In addition, pre-print BioPlex now provides a first draft of the interactome in HCT116 cells (Huttlin *et al*, 2020).

The experimental strategy behind BioPlex is based on the expression of each protein-coding open reading frame (ORF) present in the human ORFeome with an epitope tag, the affinity purification of the corresponding protein, and the confident identification of its specific protein interactors by mass spectrometry. The identification of peptides and proteins in each protein complex is performed using the Uniprot database. Hence, only proteins and alternative splicing-derived protein isoforms annotated in the Uniprot database can be detected. Using this common approach, the human interactome is necessarily made up of proteins already annotated in the Uniprot database, precluding the detection of novel unannotated proteins. Yet, beyond isoform derived proteomic diversity, multiple recent discoveries point to a general phenomenon of translation events of non-canonical ORFs in both eukaryotes and prokaryotes, including small ORFs and alternative ORFs (altORFs) (Brunet *et al*, 2020b; Orr *et al*, 2020; (Olexiouk *et al*, 2018)). Typically, small ORFs are between 10 and 100 codons, while altORFs can be larger than 100 codons. Here, we use the term altORFs for non-canonical ORFs independently of their size. On average, altORFs are ten times shorter than conventional annotated ORFs but several thousands are longer than 100 codons (Samandi *et al*, 2017). AltORFs encode alternative proteins (altProts) and are found both upstream (i.e. 5’UTR) and downstream (i.e. 3’UTR) of the reference coding sequence as well as overlapping the reference coding sequence in a shifted reading frame within mRNAs (Fig 1A-B). Additionally, RNAs transcribed from long non-coding RNA genes and pseudogenes are systematically annotated as non-coding RNAs (ncRNAs); yet, they may also harbor altORFs and encode alternative proteins (Samandi *et al*, 2017). Consequently, the fraction of multi-coding or polycistronic human genes and of protein-coding “pseudogenes” may have been largely underestimated. AltORFs translation events are experimentally detected by ribosome profiling (Orr *et al*, 2020), a method that detects initiating and/or elongating ribosomes at the transcriptome wide level (Ingolia *et al*, 2019). Alternatively, large-scale mass spectrometry detection of alternative proteins requires first the annotation of altORFs and then *in-silico* translation of these altORFs to generate customized protein databases containing the sequences of the corresponding proteins (Delcourt *et al*, 2017). This integrative approach, termed proteogenomics, has emerged as a new research field essential to better capture the coding potential and the diversity of the proteome (Nesvizhskii, 2014; Ruggles *et al*, 2017).

**Figure 1.**
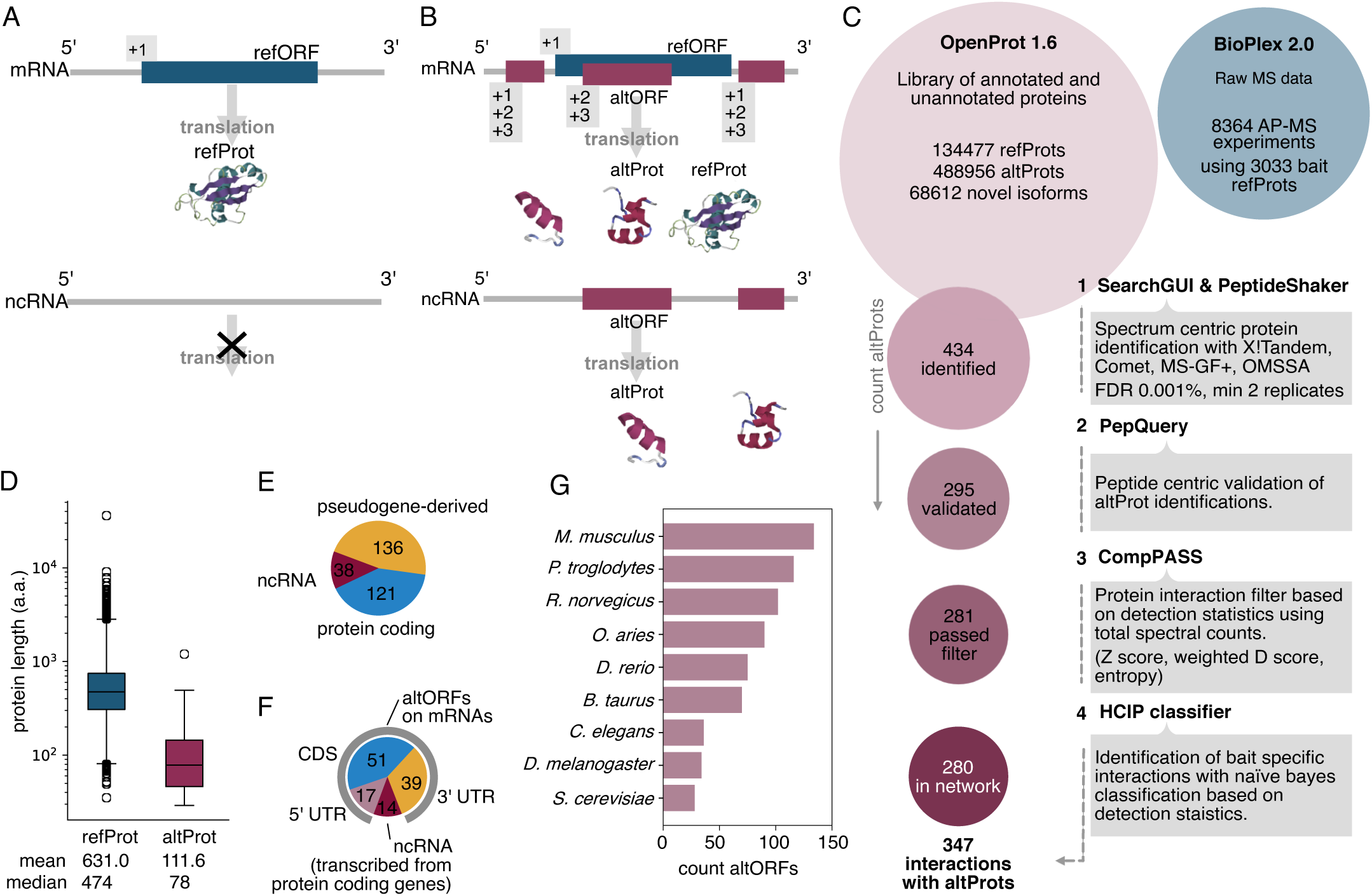
***Analysis overview and identification of alternative proteins in the human interactome.*** **A-B** The classical model of RNA transcript coding sequence annotation includes only one reference open reading frame (ORF) on mRNAs encoding a reference protein (refProt) and no functional ORF within ncRNAs (A), while the alternative translation model considers multiple proteins encoded in different reading frames in the same transcript including refProts and alternative proteins (altProt)(B). **C** Our re-analysis pipeline of high throughput AP-MS experiments from BioPlex 2.0 employs stringent criteria to ensure confident identification of both protein detection and interaction detection. Of the 434 altProts initially identified in the dataset, 280 joined the network of protein interactions after filtration. **D** AltProts are in general shorter than reference proteins. Boxes represent the inter quartile range (IQR) marked at the median and the whiskers are set at 1.5*IQR over and under the 25th and 75th percentiles. **E** Identified altProts (295) were encoded by transcripts (455) of a variety of biotypes. 121 of identified altProts are encoded by transcripts of protein coding biotype, 136 by transcripts of pseudogenes, and 38 exclusively by transcripts of non-coding biotype (ncRNA). **F** AltORFs found encoded by transcripts from genes of protein coding biotype are most often overlapping the canonical CDS or localized downstream in the 3’UTR. A significant fraction of altORFs also localize in ncRNAs of protein coding genes. CDS: coding region, UTR: untranslated region (non-coding). **G** Orthology data across 10 species from OpenProt 1.6 for detected altProts.

The translation of altORFs genuinely expands the proteome, and proteogenomics approaches using customized protein databases allows for routine MS-based detection of altProts (Brunet *et al*, 2019; Delcourt *et al*, 2018). In order to uncover altProts otherwise undetectable using the UniProt database we re-analyzed the raw MS-data from the BioPlex 2.0 interactome with our OpenProt proteogenomics database.

OpenProt contains the sequences of proteins predicted to be encoded by all ORFs larger than 30 codons in the human transcriptome. This large ORFeome includes ORFs encoding proteins annotated by NCBI RefSeq, Ensembl and Uniprot, termed here reference proteins or refProts. It also includes still unannotated ORFs that encode novel isoforms sharing a high degree of similarity with refProts from the same gene. Finally, the third category of ORFs, termed altORFs, potentially encode altProts and shares no significant sequence similarity with a refProt from the same gene (Table 1). OpenProt is not limited by the three main assumptions that shape current annotations: (1) a single functional ORF in each mRNA, typically the longest ORF; (2) RNAs with ORFs shorter than 100 codons are typically annotated as ncRNAs; and (3) RNAs transcribed from genes annotated as pseudogenes are automatically annotated as ncRNAs. Thus, in addition to proteins present in NCBI RefSeq, Ensembl and Uniprot, OpenProt also contains the sequence for novel proteins, including novel isoforms and alternative proteins (Brunet *et al*, 2019, 2020c).

**Table 1.**
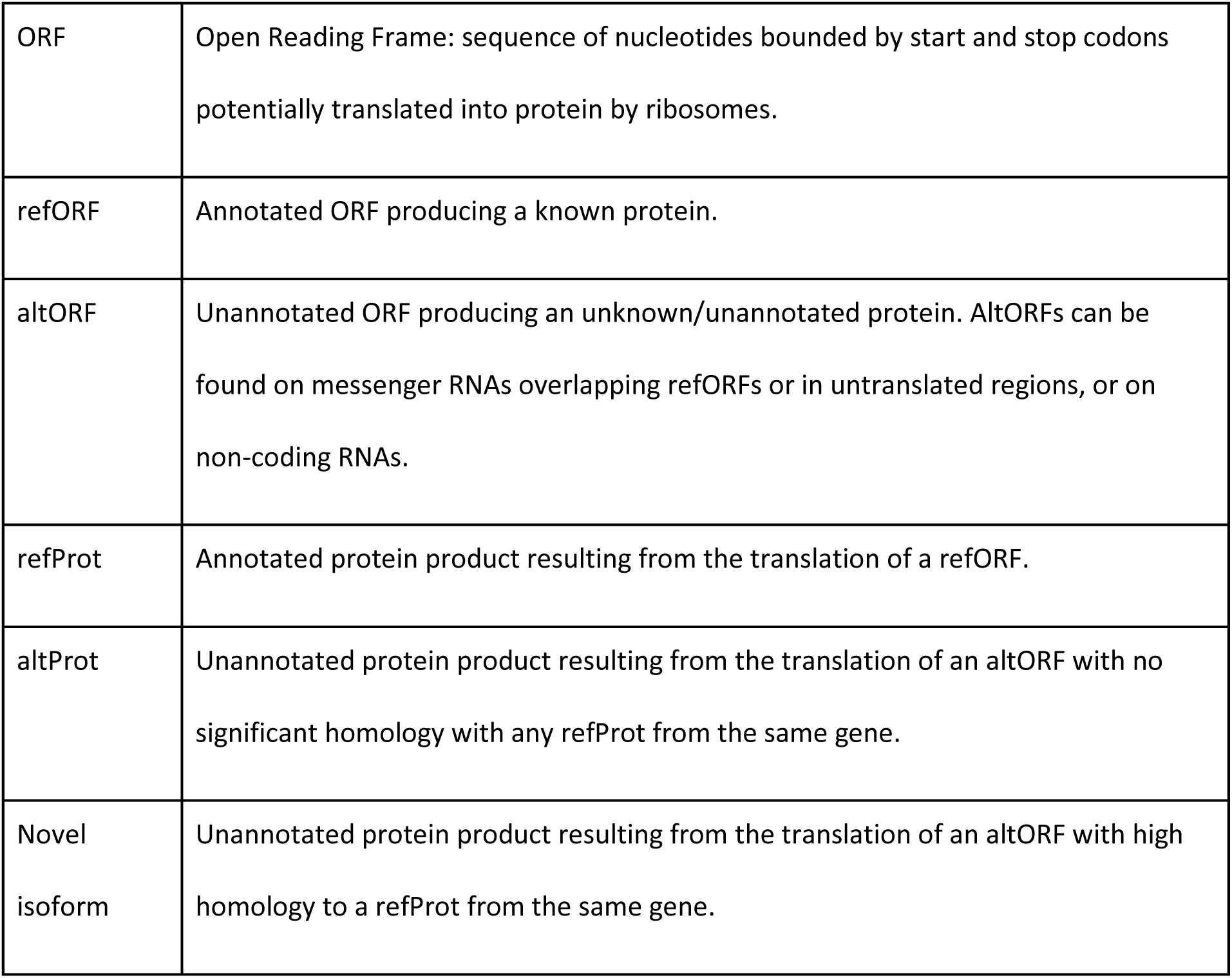
Terminology definitions

|  |  |
| --- | --- |
| ORF | Open Reading Frame: sequence of nucleotides bounded by start and stop codons potentially translated into protein by ribosomes. |
| refORF | Annotated ORF producing a known protein. |
| altORF | Unannotated ORF producing an unknown/unannotated protein. AltORFs can be found on messenger RNAs overlapping refORFs or in untranslated regions, or on non-coding RNAs. |
| refProt | Annotated protein product resulting from the translation of a refORF. |
| altProt | Unannotated protein product resulting from the translation of an altORF with no significant homology with any refProt from the same gene. |
| Novel isoform | Unannotated protein product resulting from the translation of an altORF with high homology to a refProt from the same gene. |

Using OpenProt, we were able to detect and map altProts within complexes of known proteins which increased protein diversity by including a higher number of small proteins. In addition, the data confirmed the significant contribution of pseudogenes to protein networks with 124 out of 280 altProts encoded by genes annotated as pseudogenes. We also detected many interacting proteins encoded either by the same gene or by a pseudogene and its corresponding parental gene. In sum, this work improves our knowledge of both the coding potential of the human transcriptome and the composition of protein communities by bringing diversity (i.e. small proteins) and inclusivity (i.e. proteins encoded in RNAs incorrectly annotated as ncRNAs) into the largest human protein-protein interaction (PPI) network to date.

## Results

### Re-analysis of BioPlex 2.0 mass spectrometry data and identification of preyed alternative proteins

We employed the OpenProt proteogenomics library in the re-analysis of high throughput AP-MS experiments from the BioPlex 2.0 network. Given the size of the OpenProt database (Fig 1C), the false discovery rate (FDR) for protein identification was adjusted from 1 % down to 0.001 % to mitigate against spurious identifications (Brunet *et al*, 2019). Such stringent FDR settings inevitably lead to fewer prey proteins identified; thus, our highly conservative methodology is likely to leave behind many false negatives. The BioPlex 2.0 network is built in a gene-centric manner in order to simplify the analysis by making abstraction of protein isoforms. In the current analysis, all refProts and their isoforms are also grouped under their respective gene, but results concerning altProts are necessarily given at the protein level.

In total, 434 unannotated proteins from 418 genes and 5669 refProts were identified in the re-analysis of raw MS data from the pull-down of 3033 refProts (baits), using a combination of multiple identification algorithms (Fig 1C). Since these identifications resulted from the re-analysis of raw MS data from BioPlex 2.0 with the OpenProt MS pipeline, we sought to determine the overlap between total sets of genes identified. RefProts from 4656 genes (or 85 % of total re-analysis results) were found in both the BioPlex 2.0 and in the present work (Fig EV1A), indicating that the re-analysis could reliably reproduce BioPlex results.

Our stringent approach in the identification of altProts included the use of PepQuery to validate protein detection using a peptide-centric approach (Wen *et al*, 2019). This tool includes a step which verified that altProt-derived peptides were supported by experimental spectra that could not be better explained by peptides from refProts with any post-translational modification. In addition, peptides were screened for isobaric substitutions in order to reject dubious peptides that could match refProts (Choong *et al*, 2017). A total of 295 altProt identifications were validated with PepQuery including 136 altProts encoded by pseudogenes (Table EV1). MS-based identification of short proteins with a minimum of 2 unique suitable tryptic peptides remains an important challenge and the majority of short proteins are typically detected with a single unique peptide (Slavoff *et al*, 2013; Ma *et al*, 2014). Of the 295 altProts validated by PepQuery (Table EV2), 63 complied with the Human Proteome Project PE1 level for proteins with strong protein-level evidence, Guidelines v3.0 (Deutsch *et al*, 2019).

As expected, detected altProts were much shorter than refProts with a median size of 78 amino acids versus 474 (Fig 1D; Table EV1). AltORFs encoding the 295 detected and PepQuery-validated altProts were distributed among 1029 transcripts (Table EV1) and in addition to the 136 pseudogenes derived altProts, 38 were exclusively encoded by genes of non-coding biotypes (Fig 1E). A third were found in transcripts already encoding a refProt (Fig 1E), indicating that the corresponding genes are in fact either bicistronic (two non-overlapping ORFs) or dual-coding (two overlapping ORFs) (Table EV1). Of the altProts encoded by transcripts from genes of protein coding biotype, most were encoded by a frame-shifted altORF overlapping the annotated coding sequence or downstream of the annotated coding sequence in the 3’UTR (Fig 1F). The remaining altORFs were encoded by 5’UTRs or by transcripts annotated as non-coding but transcribed from those genes of protein coding biotype. From the localization of altORFs relative to the canonical CDS in the 107 mRNA from protein coding genes we conclude that 56 of those genes are in fact bicistronic and 51 are dual-coding (Table EV1). In addition, transcripts from 7 pseudogenes have been found to encode two altProts suggesting that 3 of them are in fact dual coding and 4 are bicistronic (Table EV1).

We collected protein orthology relationships from 10 species computed by OpenProt (Fig 1G). Although 100 altProts were specific to humans, a large number had orthologs in the mouse and chimpanzee, and 28 were even conserved through evolution since yeast. 167 altProts displayed at least one functional domain signature (InterProScan, version 5.14-53.0, (Mitchell *et al*, 2019)), further supporting their functionality (Table EV1).

### Network assembly

After identification of prey proteins, CompPASS was used to compute semi-quantitative statistics based on peptide-spectral matches across technical replicates (Sowa *et al*, 2009). These metrics allow filtration of background and spurious interactions from the raw identifications of prey proteins to obtain high confidence interacting proteins (HCIP). To mitigate against the otherwise noisy nature of fast-paced high throughput approaches and to filter prey identifications down to the most confident interactions, we applied a Naïve Bayes classifier similar to CompPASS Plus (Huttlin *et al*, 2015). The classifier used representations of bait-prey pairs computed from detection statistics and assembled into a vector of 9 features as described by (Huttlin *et al*, 2015). High confidence interactions reported by BioPlex 2.0 served as target labels. HCIP classification resulted in the retention of 3.6 % of the starting set of bait-prey pairs identified (Fig EV1C). Notably, 815 baits from the original dataset were excluded after filtration because no confident interaction could be distinguished from background.

Following protein identifications and background filtration, the network was assembled by integrating all bait-prey interactions into one network (Fig 2A). All refProts and their isoforms were grouped under their respective gene, similar to the BioPlex analysis, but separate nodes are shown for altProts. In total, the re-analysis with OpenProt found 5650 prey proteins from the purification of 2218 bait proteins altogether engaged in 14029 interactions, the majority (59.1 %) of which were also reported by BioPlex 2.0 (Fig 2B). The average number of interactions per bait was 7.1. Among prey proteins, 280 altProts were found engaged in 347 interactions with 292 bait proteins.

**Figure 2.**
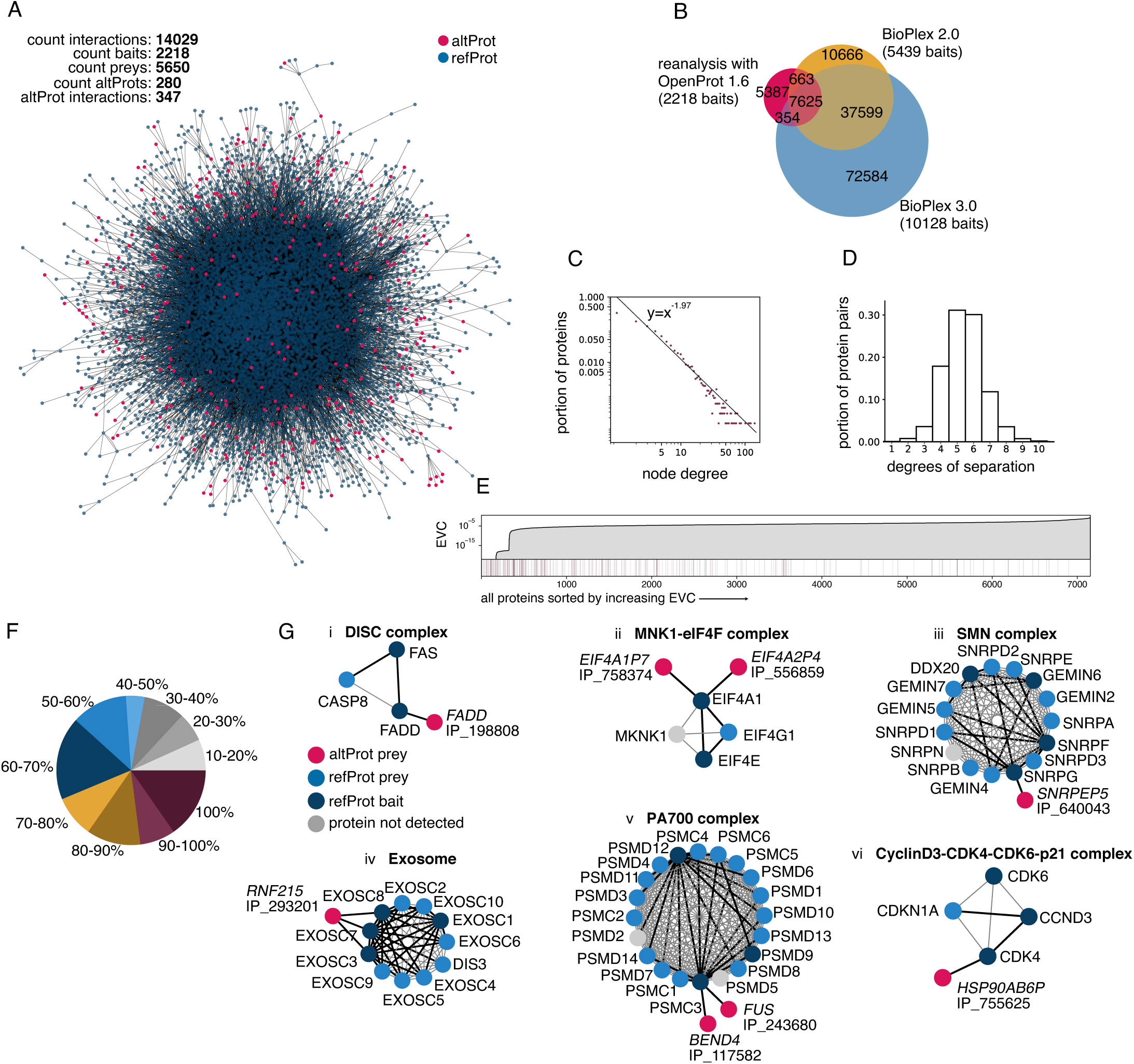
***Interaction mapping and network features of protein-protein interactions.*** **A** The largest component of the network assembled from the OpenProt based re-analysis of high throughput affinity purification mass spectrometry data from BioPlex 2.0. **B** A venn diagram of bait-prey interactions identified with the OpenProt derived re-analysis, BioPlex 2.0, and BioPlex 3.0 shows a significant overlap despite the smaller overall size of the re-analysis results (due to stringent filtration). It should also be noted that alternative proteins were not present in the BioPlex 2.0 analytical pipeline which accounts for part of the gap in overlap. **C** The degree distribution (distribution of node connectivity) follows a power law as demonstrated by a discrete maximum likelihood estimator fit. The great majority of proteins have a small number of connections while a few are highly connected (often called hubs). **D** The distribution of degrees of separation between all protein pairs (i.e. the length of the shortest path between all pairs of proteins) indicates that the network fits small-world characteristics. **E** Alternative proteins were found diffusely throughout the network and across the spectrum of eigenvector centrality (EVC) (dark lines). EVC is a relative score that indicates the degree of influence of nodes on the network; here, altProts display involvement in both influential and peripheral regions. **F** Known protein complexes from the CORUM 3.0 resource (Giurgiu *et al*, 2019) were mapped onto the network. Subunit recovery rate confirms the overall validity of the interactions confidently identified by the pipeline. All CORUM core complexes for which at least two subunits appear as baits in the network were considered. **G** Selected CORUM complexes are shown with the addition of altProts found in the interaction network of baited subunits. Black edges indicate detection in the re-analysis, grey edges indicate those only reported by CORUM.

Compared to BioPlex 2.0, a smaller total number of protein identification was expected because the OpenProt MS analysis pipeline is more stringent with a tolerance of 20 ppm on peak positions rather than 50 ppm and a 0.001 % protein FDR as opposed to 1 %. Indeed, we identified 14029 interactions in our reanalysis, compared to 56553 interactions reported by BioPlex 2.0 (Fig 2B). Among the 14029 interactions, 8288 (59.1 %) were also reported by BioPlex 2.0, and 7979 (56.8 %) were reported in the recently released (but not yet peer reviewed) BioPlex 3.0 (Fig 2B). Interestingly, 11329 interactions (20 %) from BioPlex 2.0 were not confirmed in BioPlex 3.0 using a larger number of protein baits, although the same experimental and computational methodologies were used (Fig 2B). This observation illustrates the challenge in the identification of protein-protein interactions with large-scale data given the relatively low signal to noise ratio in AP-MS data.

### Network structural features and alternative protein integration

Network theoretic analysis confirmed that the OpenProt-derived network displayed the expected characteristics of natural networks. Variability in the number of interacting partners of a given protein in a network (node degree) is typically very wide and the degree distribution that characterizes this variation follows a power-law (Bianconi & Barabási, 2001). Similar to other protein networks, the degree distribution of the OpenProt-derived network also fitted a power-law, an indication that the vast majority of proteins have few connections and a minor fraction is highly connected (also called hubs) (Fig 2C). The degree of connectivity of altProts varied between 1 and 7 whereas that of refProt was between 1 and 84. On the one hand, since long and multidomain proteins are over-represented among hub proteins (Ekman *et al*, 2006), this difference may be explained by the fact that altProts in the network were on average 6 times shorter than refProts (Fig 1D). On the other hand, none of the altProts were used as baits which also explains their lower observed connectivity since average degree was 2.5 for preys but 7.1 for baits.

The mean degrees of separation between any two proteins in the OpenProt-derived network was 5 (Fig 2D), in agreement with the small-world effect that characterizes biological networks (Wagner & Fell, 2001).

Centrality analysis allows us to sort proteins according to their relative influence on network behaviour where the most central proteins tend to be involved in the most essential cellular processes (Jeong *et al*, 2001). Here, the eigenvector centrality measure indicates that altProts are found both at the network periphery connected to refProts of lesser influence as well as connected to central refProts of high influence (Fig 2E). Since no altProts were used as baits, they are likely artificially pushed towards the edges of the network.

Known complexes from the CORUM database were mapped onto the network to assess the portion of complex subunits identified in the re-analysis (Table EV3). In most cases a majority were recovered (75 % of complexes showed ≥50 % recovery) (Fig 2F). We observed 50 altProts in the neighborhood of CORUM complex subunits that served as bait, i.e. directly interacting with the CORUM complex. Here multiple interesting patterns of altProt interactions were already noticeable: (1) altProts detected in the interactome of their respective refProts (Fig 2Gi), (2) altProts originating from pseudogenes and detected in the interactome of refProts encoded by the parental gene (Fig 2Gii-iii) and (3) altProts from protein coding genes or pseudogenes detected in network regions outside the immediate neighborhood of the related protein/gene (Fig 2Giv-vi).

The OpenProt-derived protein-protein interaction network displayed with a degree sorted circle layout showed that preyed altProts generally had a lower degree of connectivity compared to refProts (Fig 3A). This might be expected in part because no altProts were used as baits in the network, but also based on the limited range of binding capacity due to their smaller size. In order to investigate the local neighborhood of altProts, subnetworks were extracted by taking nodes within shortest path length of 2 and all edges between these for each altProt (here called second neighborhood). Notable altProts with high degree include OpenProt accessions IP_117582, a novel protein encoded by an altORF overlapping the reference coding sequence in the *BEND4* gene (Fig 3Ai), and IP_711679, encoded in a transcript of the *SLC38A10* gene currently annotated as a ncRNA (Fig 3Aii). Although these two altProts would not qualify as hub proteins per say, they seem to participate in the bridging of hubs from otherwise relatively isolated regions. Several other examples of altProts encoded by a lncRNA gene (Fig 3Aiii), in pseudogenes (Fig 3Aiv, v, vii, viii), and in protein-coding genes (Fig 3Avi, ix) integrate the network with a variety of topologies. One of these subnetworks features IP_710744, a recently discovered altProt and polyubiquitin precursor with 3 ubiquitin variants, encoded in the *UBBP4* pseudogene (Dubois *et al*, 2020). The ubiquitin variant Ubbp4^A2^ differs from canonical ubiquitin by one amino acid(T55S) and can be attached to target proteins (Dubois *et al*, 2020). Before network assembly this variant was identified reproducibly (across technical replicates) in the purification of 11 baits. Following HCIP identifications, only 3 interactions remained (Fig 3Aiv), likely because widespread identifications lead the Naïve Bayes classifier to assume non-specificity for those showing lower abundance. The 3 interactors include 2 ubiquitin ligases (*UBE2E2* (Q96LR5) and *UBE2E3* (Q969T4)) and *USP48* (Q86UV5), a peptidase involved in the processing of ubiquitin precursors.

**Figure 3.**
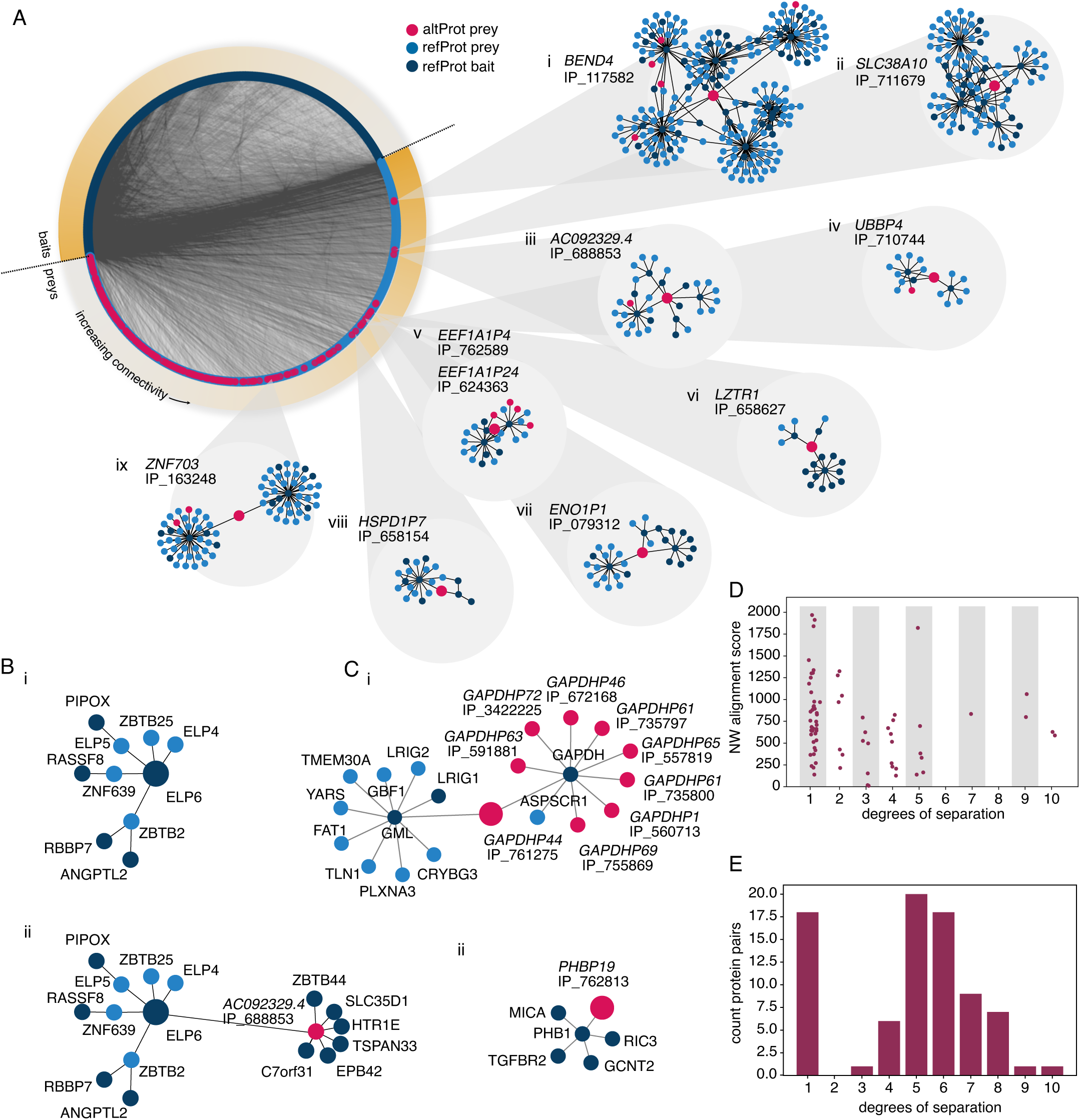
***Specific features of protein-protein interactions involving preyed alternative proteins.*** **A** Degree-sorted circular layout of the OpenProt derived full network separated by bait and preys. Direct neighbors and neighbors of neighbors (here called second neighborhood) were extracted for each altProt. Second neighborhoods of alternative proteins display a variety of topologies with some acting as bridges (iv, vi,vii,ix) and others embedded in interconnected regions (i-iii, v). Larger nodes represent the proteins for which the second neighborhood was extracted. **B** Second neighborhood of the refProt ELP6 extracted from the network assembled without altProts (i) and with altProts (ii). Inclusion of altProts in the network revealed that ELP6 connects to 6 additional proteins through its interaction with altProt IP_688853. Larger nodes represent the proteins for which the second neighborhood was extracted. **C** Detailed second neighborhood of two pseudogene-encoded altProts. (i) GAPDH refProt shows 9 altProt interactors encoded by pseudogenes of GAPDH. (ii) AltProt encoded by *PHBP19* seen in the neighborhood of the PHB refProt. Larger nodes represent the proteins for which the second neighborhood was extracted. **D** AltProt found in the direct interactome of corresponding refProt from parent genes display a wide array of sequence similarity to the refProt. Pairs of altProt-refProt from pairs of pseudogene-parental genes are slightly closer in the network if their Needleman-Wunch (NW) protein sequence global alignment score is higher. **E** The distribution of degrees of separation between altProt-refProt pairs of the same gene is bimodal with a sub-population (75 %) following a distribution similar to the full network (see Figure 2D), and the other placing altProts in the direct neighborhood of refProts from the same gene.

After observing second neighborhoods of altProts we sought to evaluate the effect of altProt inclusion into local neighborhoods of refProts. To do so we computed the eigenvector centrality of each refProt within their own second neighborhood extracted from the assembled network with and without altProts. This analysis highlighted *ELP6* which undergoes a marked reduction in eigenvector centrality in its second neighbourhood (0.67 versus 0.56) when the altProt IP_688853 (encoded by the ‘non-coding’ gene AC092329.4) is included (Fig 3Bi,ii). This shows that node influence in this region of the network is poorly understood and that identifications of novel interactors may shed light over the recent association of this gene with tumorigenesis (Close *et al*, 2012).

In total, 45 pseudogene-encoded altProts were uncovered in the direct interactome of refProts from their respective parental genes (Table EV4, shortest path length of 1), of which 2 more examples are illustrated with more details in Fig 3C.

*GAPDH* is known to have a large number of pseudogenes (Liu *et al*, 2009). Yet protein products originating from 9 *GAPDH* pseudogenes were confidently identified in the purification of the canonical GAPDH protein (Fig 3Ci). Since the glycolytic active form of this enzyme is a tetramer, we conjecture that GAPDH tetramers may assemble from a heterogenous mixture of protein products from the parental gene and many of its pseudogenes. GAPDH is a multifunctional protein (Tristan *et al*, 2011); although different posttranslational modifications may explain in part how this protein switches function (Colell *et al*, 2009), it is possible that heterologous and homologous complexes contribute to GAPDH functional diversity. Especially given that 4 of the smallest protein products from *GAPDH* pseudogenes only contain the GAPDH NAD binding domain (IPR020828; IP_735797, IP_761275, IP_735800, IP_591881), the protein encoded by *GAPDHP1* only contains the GAPDH catalytic domain (IPR020829; IP_560713), while the largest proteins from *GAPDH* pseudogenes contain both domains (IP_557819, IP_672168, IP_3422225, IP_755869) (Table EV1). The *PHB1* subnetwork highlights an interaction between *PHB1* and *PHBP19*, one of the 21 *PHB* pseudogenes (Fig 3Bii). *PHB1 and PHB2* are paralogs and the proteins they encode, PHB1 and PHB2, heterodimerize; similar to GAPDH, the PHB1/PHB2 complex is multifunctional (Osman *et al*, 2009), and the dimerization of PHB1 or PHB2 with *PHBP19*-derived IP_762813, which also contains a prohibitin domain (IPR000163), may regulate the various activities of the complex.

We reasoned that pseudogene-derived altProts directly interacting with their parental gene-derived refProts (parental protein) may result from the generally high degree of sequence similarity, particularly for refProts known to multimerize. However, although a slight reduction of alignment scores was observed with an increase in degrees of separation, the 45 altProts directly interacting with parental protein display a large variety of sequence alignment scores (Fig 3D). This suggests that direct interactions between pseudogene-derived altProts and their respective parental refProts involve other mechanisms in addition to sequence identity. Since 42 of the 45 altProts share between 1 and 7 InterPro entries with their respective parental proteins (Table EV4), protein domains may be an important mechanism driving these interactions.

The mean degrees of separation between a refProt and an altProt encoded in the same gene reveals two types of relationships (Fig 3E). 25 % (18) of altProt-refProt pairs have a degree of separation of 1, that is to say these altProts were found in the direct interactome of the corresponding refProt from the same gene. Hence, these protein pairs encoded by the same genes are clearly involved in the same function through direct or indirect physical contacts. Interestingly, 15 of these 18 altProts are encoded by dual-coding genes, i.e. with altORFs overlapping annotated CDSs. 75 % of altProt-refProt pairs follow a distribution of degrees of separation similar to the whole network (compare Fig 3E and 2D). This suggests that they are not more closely related than any other 2 proteins in the network despite shared transcriptional regulation.

### Cluster detection reveals altProts as new participants in known protein communities

Biological networks are organised in a hierarchy of interconnected subnetworks called clusters or communities. To identify these communities, unsupervised Markov clustering (MCL) (Enright *et al*, 2002) was used similarly to methodology applied to BioPlex 2.0 (Huttlin *et al*, 2017). Partitioning of the network resulted in 1045 protein clusters, 163 of which contained at least one altProt (Fig 4A). The size of altProts in these communities varied between 29 to 269 amino acids indicating that protein length may not be a limiting factor in their involvement in functional groups. Links between clusters were drawn where the number of connections between members of cluster pairs was higher than expected (detailed in Materials and Methods).

**Figure 4.**
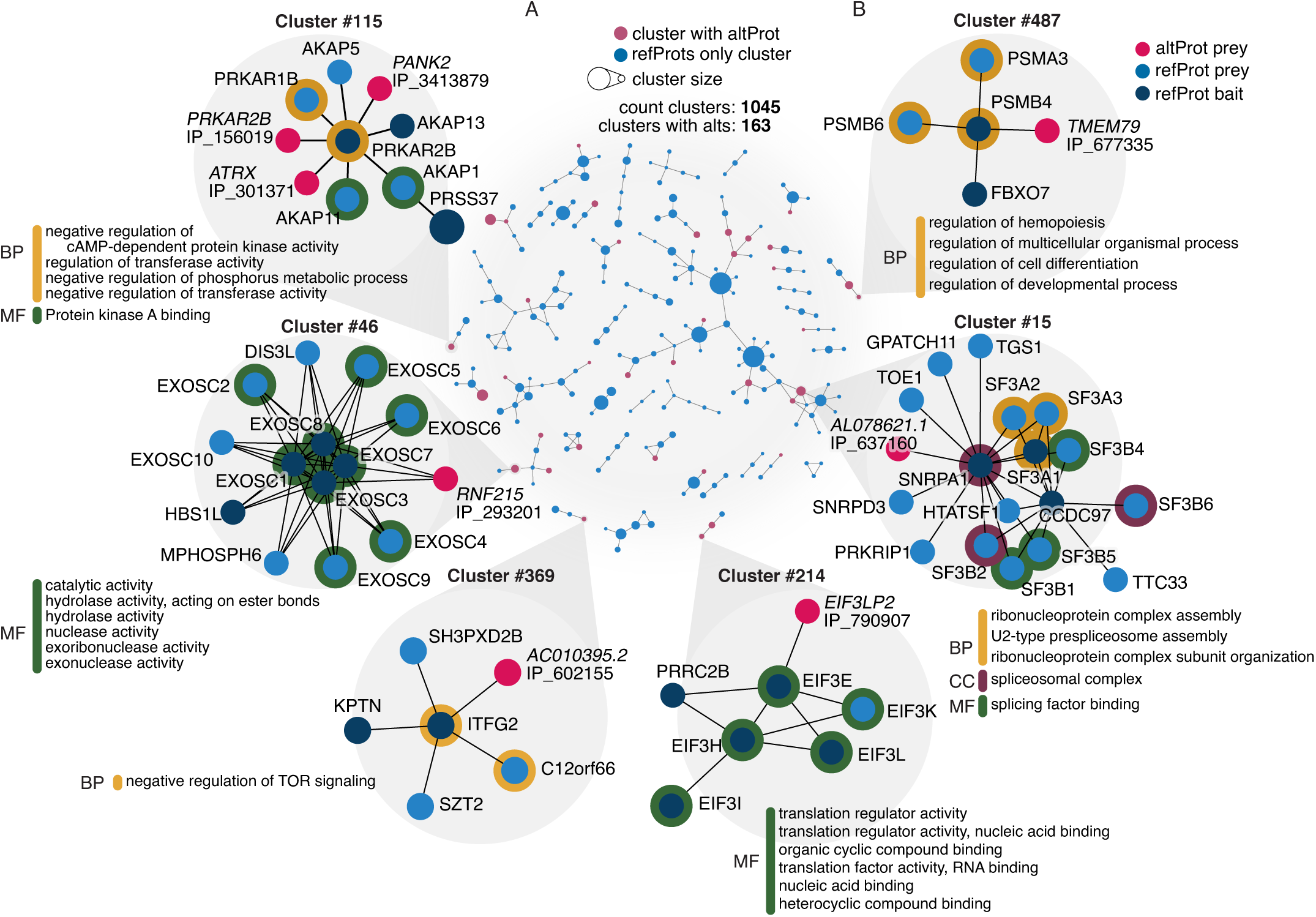
***Protein communities obtained via unsupervised community detection reveal new members*** **A** Protein communities identified via the Markov clustering algorithm (Enright *et al*, 2002). A total of 1045 clusters and 266 connections between them were identified; however, here are shown only components of 3 clusters or more for brevity. Nodes represent protein clusters sized relative to the number of proteins. Connections between clusters were determined by calculating enrichment of links between proteins in pairs of clusters using a hypergeometric test with maximal alpha value of 0.05 and correction for multiple testing was applied with 1 % FDR. **B** Focus on selected clusters showing significant enrichment of gene ontology terms. Enrichment was computed against background of whole genome with alpha value set to <0.05 Benjamini-Hochberg corrected FDR of 1 %. BP: biological process, MF: molecular function, CC: cellular compartment.

In order to assign biological function to these clusters, and therefore generate testable hypotheses about the function of altProts detected among them, enrichment of gene ontology (GO) terms was computed for each community against the background of all human genes. Several communities of different sizes showing significant GO term enrichment are detailed in Fig 4B.

45 % of identified clusters showed GO term enrichment. The same analysis with the original BioPlex network showed 57 % of clusters with GO term enrichment; possibly because a higher number of protein identifications yielded a larger network and therefore a higher probability of significant enrichment.

The altProt IP_293201 from the gene *RNF215* was identified as a novel interactor of three subunits of the RNA exosome multisubunit complex (cluster #46), suggesting a possible role in RNA homeostasis. Clusters #214 and #369 included protein communities with essential activities: the large eukaryotic initiation factor EIF3 and the recently discovered KICSTOR complex, a lysosome-associated negative regulator of mTORC1 signaling (Wolfson *et al*, 2017, 1). At least one pseudogene encoded altProt was detected in each of these clusters. Intriguingly, altProts IP_790907 (cluster #214) and IP_602155 (cluster #369) interact with the parental proteins EIF3E and ITFG2, respectively. These altProts may either compete with the parental proteins to change the activity of the complexes, or function as additional subunits since each contains a relevant functional domain (initiation factor domain, IPR019382, and ITFG2 domain, PF15907, respectively). Several subunits of the spliceosome are present in cluster #15, a protein community that includes IP_637160, a novel interactor of SNRPA1, which contains a U2A’/phosphoprotein 32 family A domain (IPR003603) where U2A’ is a protein required for the spliceosome assembly (Caspary & Séraphin, 1998). Cluster #115 contains the two regulatory subunits of PKA, PRKAR1B and PRKAR2B, which form a dimer, and several A-kinase scaffold proteins that anchor this dimer to different subcellular compartments (Di Benedetto *et al*, 2008). Three altProts interacting with PRKAR2B are also present in this cluster. Interestingly, altProt IP_156019 is encoded by an altORF overlapping the canonical PRKAR2B coding sequence; hence, *PRKAR2B* is a dual-coding gene with both proteins, the refProt and the altProt, interacting with each other. The discovery of new altProts in known protein communities demonstrates a potential for the increase in our knowledge of biological complexes.

### Disease association

The curated list of disease-gene associations published by DisGeNET relates 6,970 genes with 8,141 diseases in 32,375 associations (Piñero *et al*, 2020). After mapping this gene-disease association network onto our network of protein communities, 804 clusters of which 116 contained at least one altProt were found in association with 3,668 diseases (Fig 5A). The 116 gene-disease associations involving at least one altProt were distributed among 22 disease classes (Fig 5B). The distribution of disease-cluster associations involving altProts among the disease classes was similar to those involving refProts. Thus, no preferential association of altProts with certain disease classes could be observed.

**Figure 5.**
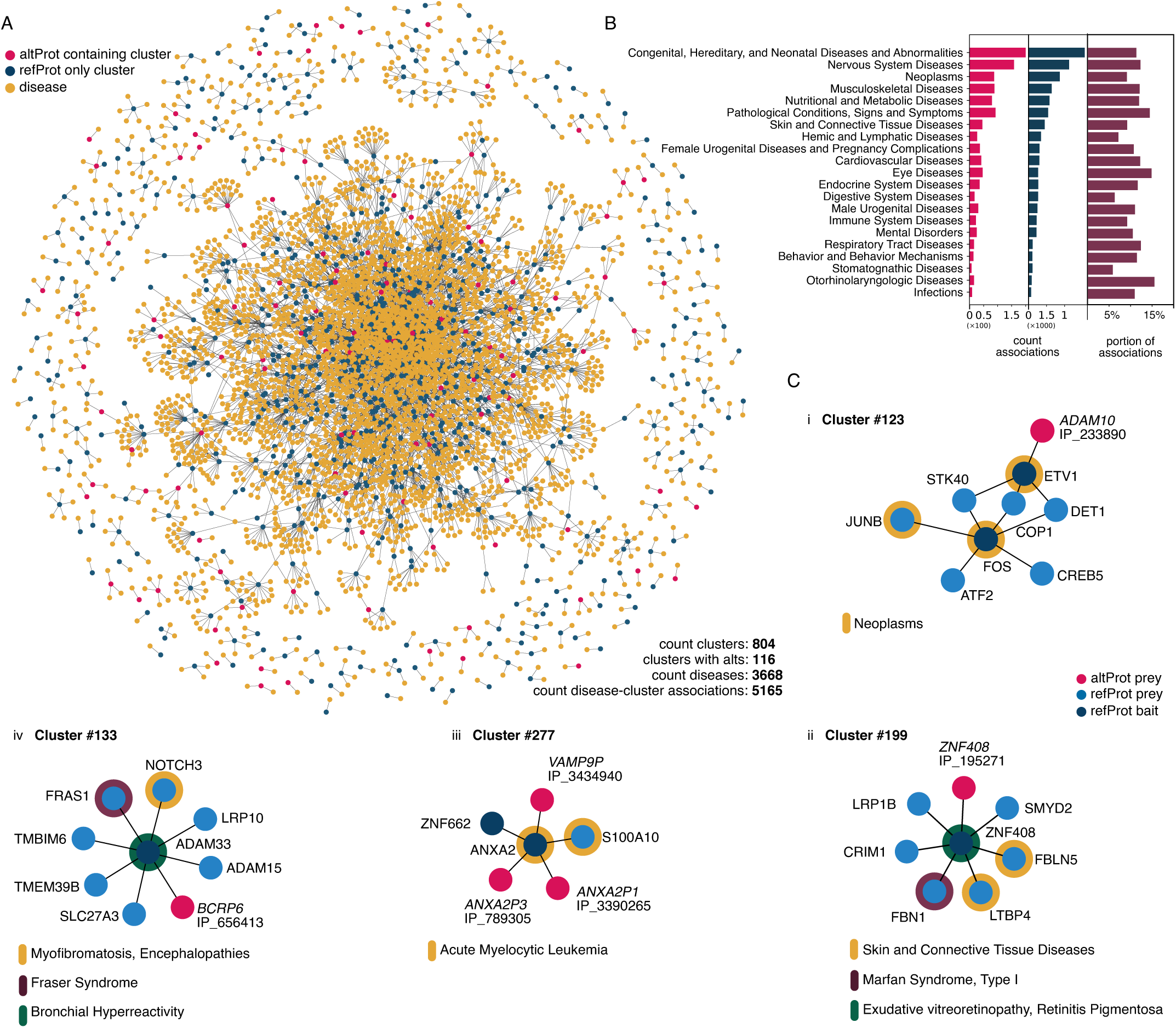
***Communities of proteins with altProt members are associated to disease phenotypes*** **A** Network of association between protein clusters (blue and red nodes) and diseases (yellow nodes) from DisGenNet. Gene-disease enrichment was computed for each pair of disease-cluster, and associations were deemed significant after hypergeometric test with alpha set to 0.01 and multiple testing correction set at maximum 1 % FDR. **B** Disease-cluster associations counted by disease classification (altProt containing clusters as red bars, and refProt only clusters as blue bars) and sorted by portion of association involving a cluster with altProts (dark red bars). **C** Focus on clusters with significant disease associations showing involvement of altProts. *ADAM10* is a gene associated with tumorigenesis and produces an altProt here detected as part of a cluster associated to neoplastic processes (i). Other cluster-disease associations include genetic connective tissue diseases involving a pair of proteins encoded by the same gene (ii) and a cluster comprising pseudogene derived altProts and parental gene refProt in association with another oncological pathology (iii). Cluster #133 (iv) highlights associations of a cluster to both rare and common diseases with a community of proteins located at the membrane.

A selection of subnetworks illustrates how altProts associate with different diseases (Fig 5C). *ADAM10* encodes a transmembrane refProt with metalloproteinase activity. Among protein substrates that are cleaved by ADAM10 and shed from cells, some act on receptors and activate signaling pathways important in normal cell physiology (Reiss & Saftig, 2009). Overexpression of this protease or increased shedding of tumorigenic proteoforms results in overactivation of signaling pathways and tumorigenesis (Murphy, 2008; Smith *et al*, 2020). IP_233890 is an altProt expressed from bicistronic *ADAM10* and its association with a subnetwork of transcription factors involved in tumorigenesis may further clarify the role of that gene in cancer (Fig 5Ci). Cluster #199 illustrates the association of a pair of refProt/altProt expressed from the same dual-coding gene, ZNF*408*, with three different diseases (Fig 5Cii). The implication of pseudogene-derived altProts is emphasized by the association of three of them with Acute Myelocytic Leukemia through their interaction with *ANXA2* (Fig 5C iii). Two of these interactions occur between a refProt from the parental gene and altProts encoded by two of its pseudogenes.

Cluster #133 relates proteins localized at the membrane with roles in intercellular signaling, development and organogenesis, as well as fatty acids transport proteins (Mahesh, 2013; Drazyk *et al*, 2019; Short *et al*, 2007, 1; Kim *et al*, 2020). AltProt IP_656413 associated with this cluster is coded by a pseudogene of the breakpoint cluster protein BCR, a Rho GTPase activating protein. IP_656413 is predicted to have a Rho GTPase activating protein domain InterProScan analysis (IPR000198) (Table EV1). Associations of this cluster with diseases both common (bronchial hypersensitivity) and rare (Fraser syndrome) highlight the potential of deeper protein coding annotations coupled with network proteomic studies to unveil novel members relevant to a wide array of pathological phenotypes. Characterization of the role of this altProt at the membrane, likely involved in intercellular signaling, may yield mechanistic insight surrounding associated pathologies.

### Functional validation of protein-protein interactions involving an alternative protein

Interactions representative of the three following classes of complexes involving altProts were selected for further experimental validation: an altProt encoded by a dual-coding gene and interacting with the respective refProt, an altProt expressed from a pseudogene and interacting with the refProt encoded by the parental gene, and an altProt interacting with a refProt coded by a different gene.

The dual-coding *FADD* gene expresses altProt IP_198808 in addition to the conventional FADD protein, and both proteins interact within the DISC complex (Fig 2Gi). We took advantage of a previous study aiming at the identification of the FADD interactome to test whether this altProt may also have been missed in this analysis because the protein database used did not contain altProt sequences (Eyckerman *et al*, 2016). In this work, the authors developed a new method called ViroTrap to isolate native protein complexes within extracellular virus-like particles to avoid artefacts of cell lysis in AP-MS. Among the baits under study FADD was selected to isolate the native FADD complex. First, we used the peptide-centric search engine PepQuery to directly test for the presence or the absence of IP_198808-derived specific peptides in the FADD complex datasets. Rather than interpreting all MS/MS spectra, this approach tests specifically for the presence of the queried peptides (Ting *et al*, 2015). Indeed, two unique peptides from IP_198808 were detected in each of the replicates of that study via PepQuery (Fig EV3A i,v). Second, we used a conventional spectrum-centric and database search analysis with the UniProt database to which was added the sequence of IP_198808. The altProt was identified in the FADD interactome (Fig EV3B) with 4 unique peptides (Fig EV3A i,iii,iv,v). In cells co-transfected with Flag-FADD and IP_198808-GFP, FADD formed large filaments (Fig 6A, right), previously labelled Death Effector Filaments (Siegel *et al*, 1998). IP_198808 co-localized in the same filaments in the nucleus, while the cytosolic filaments contained FADD only. Finally, this interaction was validated by co-immunoprecipitation (Fig 6A, left). These proteomics, microscopic and biochemical approaches confirmed the interaction between the two proteins encoded in dual-coding *FADD*.

**Figure 6.**
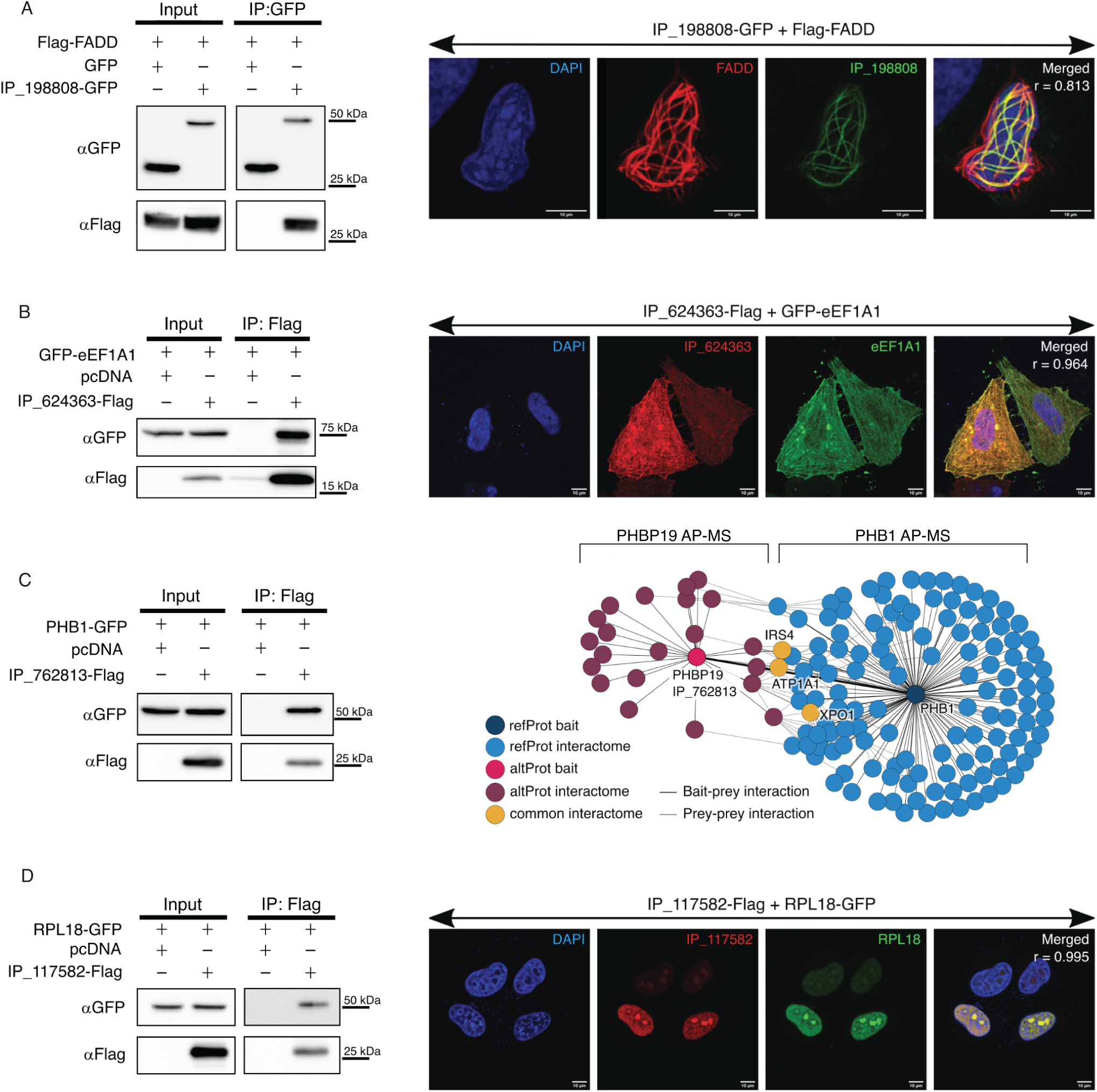
***Experimental validation of refProt-altProt interactions.*** **A** Validation of FADD and IP_198808 protein interaction encoded by a bicistronic gene. Left panel: Immunoblot of co-immunoprecipitation with GFP-trap sepharose beads performed on HEK293 lysates co-expressing Flag-FADD and IP_198808-GFP or GFP only. Right panel: confocal microscopy of HeLa cells co-transfected with IP_198808-GFP (green channel) and Flag-FADD construct immunostained with anti-Flag (red channel). r = Pearson’s correlation. The associated Manders’ Overlap Coefficients are respectively M1= 0.639 and M2 = 0.931. **B** Validation of eEF1A1 and IP_624363 protein interaction encoded from a pseudogene/parental gene couple. Left panel: immunoblot of co-immunoprecipitation with Anti-FLAG magnetic beads performed on HEK293 lysates co-expressing GFP-eEF1A1 and IP_624363-Flag or pcDNA3.1 empty vector with IP_624363-Flag constructs. Right panel: confocal microscopy of HeLa cells co-transfected with GFP-eEF1A1 (green channel) and IP_624363-Flag constructs immunostained with anti-Flag (red channel). r = Pearson’s correlation. The associated Manders’ Overlap Coefficients are respectively M1= 0.814 and M2 = 0.954. **C** Validation of PHB1 and IP_762813 protein interaction encoded by a pseudogene/parental gene couple. Left panel: immunoblot of co-immunoprecipitation with Anti-FLAG magnetic beads performed on HEK293 lysates co-expressing PHB1-GFP and IP_762813-Flag or pcDNA3.1 empty vector with IP_762813-Flag constructs. Right panel: Comparison of the interaction network of IP_762813-Flag (purple) and PHB1-GFP (blue) from independent affinity purification mass spectrometry (AP-MS) of both proteins. 3 independent AP-MS for each protein. **D** Validation of RPL18 and IP_117582 protein interaction. Left panel: immunoblot of co-immunoprecipitation with Anti-FLAG magnetic beads performed on HEK293 lysates co-expressing RPL18-GFP and IP_117582-Flag or pcDNA3.1 empty vector with IP_117582-Flag constructs. Right panel: confocal microscopy of HeLa cells co-transfected with RPL18-GFP (green channel) and IP_117582-Flag constructs immunostained with anti-Flag (red channel). r = Pearson’s correlation. The associated Manders’ Overlap Coefficients are respectively M1= 0.993 and M2 = 0.972. All western blots and confocal images are representative of at least 3 independent experiments.

Next, we selected 2 pairs of interactions of an altProt expressed from a pseudogene with a refProt expressed from the corresponding parental gene. The interaction between altProt IP_624363 encoded in the *EEF1AP24* pseudogene and EEF1A1 (Fig 3Av) was confirmed by co-immunoprecipitation from cell lysate from cells co-transfected with GFP-eEF1A1 and IP_624363 (Fig 6B, left). Both proteins also displayed strong co-localization signals (Fig 6B, right). In order to validate the interaction between *PHBP19*-encoded IP_762813 and PHB1, we performed two experiments. First, PHB1 co-immunoprecipitated with IP_762813 using cell lysates from cells co-transfected with PHB1-GFP and IP_762813-Flag (Fig 6C, left). Second, we performed independent AP-MS experiments for both IP_762813 and PHB1 in HEK293 cells. We confirmed the presence of PHB1 in the interactome of IP_762813 and the presence of IP_762813 in the interactome of PHB1 (Fig 6D, right). Interestingly, we observed shared interactors between IP_762813 and PHB1 (IRS4 (O14654), ATP1A1 (P05023) and XPO1 (O14980)), as well as interactors specific to each. Prey-prey interactions from STRING also showed a certain interconnectivity of both interactomes, whilst each retained unique interactors (Fig EV3C).

The altProt IP_117582 encoded in the *BEND4* gene is one of the most central and most connected alternative proteins in our network (Fig 3A). The interaction with RPL18 was tested and confirmed by co-immunoprecipitation in cells co-transfected with RPL18-GFP and IP_117582-Flag (Fig 6D, left), and their co-localization was also confirmed by immunofluorescence (Fig 6D, right).

## Discussion

The discovery of unannotated altProts encoded by ORFs localized in “non-coding” regions of the transcriptome raises the question of the function of these proteins. The translation of altProts may result from biological translational noise producing non-bioactive molecules. Alternatively, altProts may play important biological roles (Orr *et al*, 2020). Here, we addressed the issue of the functionality of altProts by testing their implication in protein-protein interactions. We have reanalyzed the Bioplex 2.0 proteo-interactomics data using the proteogenomics resource OpenProt which provides customized databases for all ORFs larger than 30 codons in 10 species (Brunet *et al*, 2019, 2020c). Under stringent conditions, a total of 295 prey altProts were detected, of which 280 could be confidently mapped in the network of 292 bait refProts. 136 altProts are expressed from pseudogenes, 121 from dual-coding and bicistronic genes, and 38 from transcripts annotated as ncRNA but should in fact be protein-coding. In addition to revealing new members of protein communities, this study lends definitive support to the functionality of hundreds of altProts and provides avenues to investigate their function.

The detection of 295 altProts under stringent conditions confirms the hindrance introduced by three assumptions of conventional annotations: (1) eukaryotic protein-coding genes are monocistronic; (2) RNAs transcribed from genes annotated as pseudogenes are ncRNAs; and (3) ncRNAs are annotated as such based on non-experimental criteria, including the largely used 100 codons minimal length (Dinger *et al*, 2008). The persistence of these assumptions in conventional genomic annotations limits the repertoire of proteins encoded by eukaryotic genomes (Brunet *et al*, 2018). It remains possible that functional altORFs in regions of the transcriptome annotated as non-coding are exceptions and that a large fraction of genes and RNAs comply with current assumptions. However, an ever-increasing number of proteogenomics studies demonstrate that thousands of altORFs and their corresponding proteins are translated (Samandi *et al*, 2017; Chen *et al*, 2020).

Conventional annotations introduce some confusion by opting to create a new gene entry within a previously annotated gene where a novel protein product has been reported or where novel transcripts have been mapped, rather than annotate a second ORF in the initial gene. The result is that some genomic regions have been assigned a second gene in the same orientation, nested within a previously annotated gene. This is the case for pseudogene *ENO1P1* (Ensembl: ENSG00000244457; genomic location: chr1: 236,483,165-236,484,468 (GRCh38.p13)) which overlaps the protein coding gene *EDARADD* (Ensembl: ENSG00000186197; genomic location: chr1:236,348,257-236,502,915 (GRCh38.p13)) which also encodes altProt IP_079312. Thus, as a result of this annotation, a pseudogene (*ENO1P1*) is nested within a protein-coding gene (*EDARADD*). Similarly, a second protein-coding gene termed *AL022312.1* (Ensembl: ENSG00000285025; genomic location: chr22: 39,504,231-39,504,443 (GRCh38.p13)) was added within the protein-coding *MIEF1* gene (Ensembl: ENSG00000100335; genomic location: chr22:39,499,432-39,518,132 (GRCh38.p13)) to annotate the recently discovered altORF upstream of the *MIEF1* CDS (Samandi *et al*, 2017; Vanderperre *et al*, 2013). We suggest that recognizing the polycistronic nature of some human genes to be able to annotate multiple protein-coding sequences in the same gene is more straightforward than annotating additional small genes nested in longer genes in order to comply with monocistronic annotations.

The involvement of 280 altProts in 347 of the 14029 protein-protein interactions in the current network (or 2.5 %) represents a sizable number of previously missing nodes and edges and contributes to the understanding of network topology. The impact of altProt inclusion on network structure is revealed by the bridging role many seem to play between interconnected regions (Fig 3Ai-ix). This linkage of otherwise independent complexes introduces major changes to network structure shown to be related to biological system state (e.g. cell type) (Huttlin *et al*, 2020). Results from the current analysis are thus anticipated to yield insight regarding molecular function and mechanisms of protein complexes in the contexts of cell type and other suborganismally defined states (Huttlin *et al*, 2020). Indeed, the presence of altProts in protein communities associated with known function and/or diseases makes it possible to generate testable hypotheses regarding their role in physiological and pathological mechanisms (Leblanc & Brunet, 2020).

An important observation stemming from the current study is that many pseudogenes encode one altProt in the network, including some encoding 2 altProts. Strikingly, several altProts expressed from pseudogenes interact with their respective parental protein. This suggests that pseudogene-encoded altProts are functional paralogs and that their incorporation into homomeric protein complexes of the parental protein could modulate or change the activity of the parental complex. Such function would be reminiscent of the role of homomers and heteromers of paralogs in the evolution of protein complexes in yeast, allowing structural and functional diversity (Marchant *et al*, 2019; Pereira-Leal *et al*, 2007). The GAPDH subnetwork with its 9 pseudogene-encoded altProts is particularly striking. Besides its canonical function in glycolysis, GAPDH displays a variety of different functions in different subcellular locations, including apoptosis, DNA repair, regulation of RNA stability, transcription, membrane fusion, and cytoskeleton dynamics (Colell *et al*, 2009; Sirover, 2012; Tristan *et al*, 2011). We propose that the incorporation of different paralog subunits in this multimeric complex results in the assembly of different heteromeric complexes and may at least in part entail such functional and localization diversity. This hypothesis is in agreement with the speculation that the diversity of functions associated with GAPDH correlates with the remarkable number of GAPDH pseudogenes (Liu *et al*, 2009).

Among the 274 genes encoding the 280 altProts inserted in the network, 18 encode refProt/altProt pairs that specifically interact with each other, which implies that these pairs are involved in the same function. Such functional cooperation between a refProt and an altProt expressed from the same eukaryotic gene confirms previous observations in humans (Samandi *et al*, 2017; Chen *et al*, 2020; Bergeron *et al*, 2013; Klemke *et al*, 2001). Dual-coding genes are common in viruses (Chirico *et al*, 2010) and proteins expressed from viral overlapping ORFs often interact (Pavesi *et al*, 2018). The general tendency of physical or functional interaction between two proteins expressed from the same gene should help decipher the role of newly discovered proteins provided that functional characterization of the known protein is available. Molecular mechanisms behind the functional cooperation of such protein pairs remain to be explored.

Furthermore, several pairs of proteins encoded by the same gene but acting in distant parts of the network have also been identified. Could these altProts be a source of cross talk between functional modules under the same regulation at the genetic level, but multiplexed at the protein function level?

The current study shows that the 280 altProts incorporated in the network differ from refProts by their size (6 times smaller in average) but do not form a particular class of gene products; rather they are members of common communities present throughout the proteomic landscape. Initial serendipitous detection of altProts subsequently called for proteogenomics approaches which widened discoveries via systematic and large-scale detection (Peeters & Menschaert, 2020; Brunet *et al*, 2020b). System resilience and biodiversity have long been linked in the ecology literature (Peterson *et al*, 1998); by analogy the increased proteomic diversity due to altProts could be a contributing factor to this effect in cellular systems. To find out the extent to which altProts play widespread and important biological functions will require more studies in functional genomics.

## Materials & Methods

### Classification of proteins, transcripts and genes

Reference proteins (RefProts) are known proteins annotated in NCBI RefSeq, Ensembl and/or UniProt. Novel isoforms are unannotated proteins with a significant sequence identity to a RefProt from the same gene; for these isoforms, BLAST search yields a bit score over 40 for an overlap over 50% of the queried reference sequence. Alternative proteins (AltProts) are unannotated proteins with no significant identity to a RefProt from the same gene.

Alternative open reading frames (altORFs) correspond to unannotated ORFs predicted to encode proteins with no significant identity to any other annotated protein.

We classify RNA transcripts as dual coding or bi-cistronic based on the relative position of the ORFs on the transcript. If they are overlapping (i.e. if they share nucleotides) we classify the transcript as dual coding, if they are sequential (i.e. share no nucleotides) we classify it as bicistronic. Gene classification with this respect is inherited from the classification of transcript it produces. Note that transcripts and genes can hold both dual coding and bicistronic classifications.

### Reanalysis of AP-MS data

Files obtained from the authors of the BioPlex 2.0 contained the results of 8,364 affinity purification-mass spectrometry (AP-MS) experiments using 3033 bait proteins (tagged with GFP) in 2 technical replicates or more barring missing replicates and corrupted files (Huttlin *et al*, 2017, 2015). Files were converted from RAW to MGF format using Proteowizard 3.0 and searched with SearchGUI 2.9.0 using an ensemble of search engines (Comet, OMSSA, X!Tandem, and MS-GF+). Search parameters were set to a precursor ion tolerance of 4.5 ppm and fragment ion tolerance of 20 ppm, trypsin digestion with a maximum of 2 missed cleavages, and variable modifications included oxidation of methionine and acetylation of N termini. The minimum and maximum length for peptides were 8 and 30 amino acids respectively. Search results were aggregated using PeptideShaker 1.13.4 with a 0.001 % protein level false discovery rate (FDR) as described previously (Brunet *et al*, 2019). In addition to already annotated proteins, the OpenProt database includes all predicted altProts and novel isoforms. Since large databases result in a large increase of false positive rates (Jeong *et al*, 2012; Nesvizhskii, 2014), this effect is balanced using an FDR of 0.001% as previously described (Brunet *et al*, 2020; Brunet & Roucou, 2019) (PMID: 32780568, 31033953). The protein library contained a non redundant list of all reference proteins from Uniprot (release 2019_03_01), Ensembl (GRCh38.95), and RefSeq (GRCh38.p12) (134477 proteins) in addition to all alternative protein (488956 proteins) and novel isoforms (68612 proteins) predictions from OpenProt 1.6. AltProt identifiers throughout the current article are accessions from OpenProt starting with “IP_”. The library was concatenated with reversed sequences for the target decoy approach to spectrum matching.

### Validation of altProt identifications

Novel protein identifications were supported by unique peptides. An additional peptide centric approach was used to validate that spectra supporting such peptides could not be better explained by peptides from refProts with post-translational modifications. PepQuery allows the search of specific peptides in spectra databases using an unrestricted modification search option (Wen *et al*, 2019). All possible peptide modifications from UniMod artifact and post translational modifications were considered when ensuring unicity of spectral matches (downloaded March 2020) (Dm & Js, 2004). AltProt sequences with peptides validated with PepQuery have been submitted to the Uniprot Knowledge Base.

### Obtaining spectral counts

Because altProts are smaller than refProts they have a lower number of uniquely identifying peptides. For this reason altProts with at least one unique peptide across multiple replicates were considered, but only refProts identified with at least two unique peptides across multiple replicates were retained for downstream analysis. Spectra shared among refProts were counted in the total spectral count of each protein. Spectra assigned to altProts were counted only if unique to the protein or shared with another altProt. Spectra shared between an altProt and at least one refProt were given to the refProt. RefProt spectral counts were combined by gene following the methodology of the original study; however, it was necessary to keep altProts separate as many are encoded by genes that already contain a refProt or other altProts.

### Interactions scoring

Following protein identifications, high confidence interacting proteins (HCIPs) were identified following the method outlined in the original study (Huttlin *et al*, 2015). Briefly, the CompPASS R package was first used to compute statistical metrics (weighted D-score, Z score, and entropy) of prey identification based on peptide spectrum match (PSM) counts. The results from CompPASS were then used to build a vector of 9 features (as described in (Huttlin *et al*, 2015)) for each candidate bait-prey pair which were passed to a Naive Bayes classifier (CompPASS Plus) tasked with the discrimination of HCIP from background identifications. The original study also included a class for wrong identification, but since decoy information was unavailable and because our approach employs a FDR three orders of magnitudes lower in the identification step, a third class was not deemed necessary. The classifier was trained in cross-validation fashion using 96 well plate batches as splits and protein-protein interactions from the original study as target labels for true interactors.

Threshold selection was implemented considering the Jaccard overlap (equation i), recall (equation ii), precision and F1 score (equation iv) metrics between networks resulting from the re-analysis and the original study. The main differences between the OpenProt derived re-analysis and BioPlex 2.0 lie in the total spectral counts resulting from the use of different search algorithms and more stringent FDR. It was thus important to tune model threshold selection to maximally reproduce results from the original study (Figure EV1B). A threshold of 0.045 was selected as it compromised well between optimal Jaccard overlap, F score, and precision (Fig EV1A).

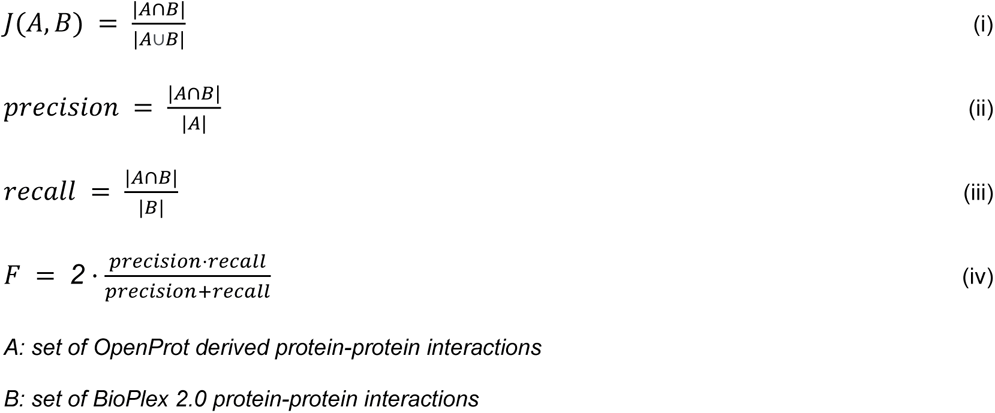

### Network assembly and structural analysis

Bait-prey pairs classified as HCIP were combined into an undirected network using genes to represent refProt nodes and OpenProt protein accessions to represent altProt nodes. The Networkx 2.5 Python package was used for network assembly and all network metrics calculations.

The power law fit to the degree distribution was computed with the discreet maximum likelihood estimator described by (Clauset *et al*, 2009).

A list of known protein complexes from CORUM 3.0 (Giurgiu *et al*, 2019) (core complexes, downloaded March 2020) was mapped onto the resulting network to assess the validity of identified interactions (Table EV3). Only complexes in which at least two subunits corresponded to baits present in the network were selected for downstream analyses. The portion of subunits identified in the direct neighbourhood of baits was computed for each complex.

### Patterns of interactions involving altProt and refProts

We aimed to assess the relationship between pseudogene-derived altProts and their corresponding refProts from parental genes, in terms of their sequence similarity and their degrees of separation in the network. Parent genes of pseudogenes were selected via the psiCUBE resource (Sisu *et al*, 2014) combined with manual curation using Ensembl. Needleman Wunch global alignment algorithm (with BLOSUM62 matrix) as implemented by the sciki-bio Python package (version 0.5.5) was used as a similarity measure between protein sequences. To assess degrees of separation, shortest path lengths were computed both for altProt-refProt pairs of pseudogene-parental gene and altProt-refProt pairs encoded by the same gene. For the former, when the refProt was not present in the network, or when no path could be computed between nodes, the shortest path length was computed using a mapping of either the BioPlex 2.0 or BIOGRID networks (Stark *et al*, 2006).

### Community detection via clustering

A Python implementation of the markov clustering (MCL) algorithm (https://github.com/GuyAllard/markov_clustering) was used to partition the network into clusters of proteins (Enright *et al*, 2002). Various values of the inflation parameter between 1.5 and 2.5 were attempted and, similarly to the original study, a value of 2.0 was selected as it compared favorably with known protein complexes. Only clusters of 3 proteins or higher were retained yielding a total of 1045 clusters. Connections between clusters were determined by calculating enrichment of links between proteins in pairs of clusters using a hypergeometric test with alpha value set to <0.05 and a Benjamini-Hochberg corrected FDR of 1 %. A total of 266 pairs of clusters were found to be significantly connected.

### Disease association

A list of 32,375 disease-gene associations curated by DisGeNET (downloaded March 2020) was mapped onto the network of 1045 protein communities. A disease was associated with a cluster when it was deemed enriched in genes associated with the disease as calculated by hypergeometric testing, with alpha value set to <0.01 Benjamini-Hochberg corrected FDR of 1 %.

### Gene Ontology Enrichment

Gene Ontology term enrichments for both altProt second neighborhoods and protein clusters were computed using the GOAtools Python package (version 1.0.2). Count propagation to parental terms was set to true, alpha value to 0.05, with a Benjamini-Hochberg corrected FDR of 1 %.

### Classification of proteins, transcripts and genes

Reference proteins (RefProts) are known proteins annotated in NCBI RefSeq, Ensembl and/or UniProt. Novel isoforms are unannotated proteins with a significant sequence identity to a RefProt from the same gene. These isoforms are identified with a BLAST search filtered for a bit score over 40 for an overlap over 50% of the queried reference sequence. Alternative proteins (AltProts) are unannotated proteins with no significant identity to a RefProt from the same gene. Importantly, altProts may share a sequence similarity with a protein from a different gene, for example in the case of pseudogene-encoded altProts and the protein derived from the parental gene.

Alternative open reading frames (altORFs) correspond to unannotated ORFs predicted to encode proteins with no significant identity to any other annotated protein.

We classify RNA transcripts as dual coding or bicistronic based on the relative position of the ORFs on the transcript. If they are overlapping (i.e. if they share nucleotides) we classify the transcript as dual coding, if they are sequential (i.e. share no nucleotides) we classify it as bicistronic. Gene classification with this respect is inherited from the classification of transcript it produces. Note that transcripts and genes can hold both dual coding and bicistronic classifications.

### Cloning and antibodies

All nucleotide sequences were generated by the Bio Basic Gene Synthesis service, except for pcDNA3-FLAG-FADD, a kind gift from Jaewhan Song (Addgene plasmid # 78802 ; http://n2t.net/addgene:78802 ; RRID:Addgene_78802). IP_117582, IP_624363, and IP_762813 were all tagged with 2 FLAG (DYKDDDDKDYKDDDDK) at their C-terminal. IP_198808 was tagged with eGFP at its C-terminal. All altProt coding sequences were subcloned into a pcDNA3.1- plasmid. The coding sequences of RPL18, eEF1A1 and PHB were derived from their canonical transcript (NM_000979.3, NM_001402.6, NM_001281496.1 respectively). RPL18 and PHB were tagged with eGFP at their C-terminal and eEF1A1 was tagged with eGFP at its N-terminal. All refProt coding sequences were subcloned into a pcDNA3.1- plasmid.

### Cell culture, transfections and immunofluorescence

HEK293 and HeLa cultured cells were routinely tested negative for mycoplasma contamination (ATCC 30–1012K). Transfections, immunofluorescence, confocal analyses were carried out as previously described (Brunet *et al*, 2020a). Briefly, transfection was carried with jetPRIME®, DNA and siRNA transfection reagents (VWR) according to the manufacturer’s protocol. To note, only

0.1 μg of pEGFP DNA versus 3 μg IP_198808-GFP was used for transfection in 100 mm petri dishes to compensate for its higher transfection and expression efficiency. Cells were fixed in 4 % paraformaldehyde for 20 mins at 4°C, solubilized in 1 % Triton for 5 mins and incubated in blocking solution (10 % NGS in PBS) for 20 mins. The primary antibodies were diluted in the blocking solution as follows: anti-Flag (Sigma, F1804) 1/1000. The secondary antibodies were diluted in the blocking solution as follows: anti-mouse Alexa 647 (Cell signaling 4410S) 1/1000. All images were taken on a Leica TCS SP8 STED 3X confocal microscope.

### Affinity Purification and western blots

Immunoprecipitation experiments via GFP-Trap (ChromoTek, Germany) were carried out as previously described (Samandi *et al*, 2017), while experiments via Anti-FLAG® M2 Magnetic Beads (M8823, Sigma) were conducted according to the manufacturer’s protocol with minor modifications. Briefly, HEK293 cells were lysed in the lysis buffer (150 mM NaCl, 50 mM Tris pH 7.5, 1 % Triton, 1 x EDTA-free Roche protease inhibitors) and incubated on ice for 30 mins prior to a double sonication at 12 % for 3 seconds each (1 min on ice between sonications). The cell lysates were centrifuged, the supernatant was isolated and the protein content was assessed using BCA assay (Pierce). Anti-FLAG beads were conditioned with the lysis buffer. 20 µL of beads were added to 1 mg of proteins at a final concentration of 1 mg/mL and incubated overnight at 4°C. Then, the beads were washed 5 times with the lysis buffer (twice with 800 µL and twice with 500µL) prior to elution in 45 µL of Laemmli buffer and boiled at 95°C for 5 min. For co-immunoproecipitation of PHB1-GFP and RPL18-GFP, stringent wash were done with modified lysis buffer (250 mM NaCl + 20 µg/ml peptide FLAG (F3290 Sigma)) prior to elution with 200µg/ml peptide FLAG. Eluates were loaded onto 10 % SDS-PAGE gels for western blotting of GFP and FLAG tagged proteins. 40 µg of input lysates were loaded into gels as inputs. Western blots were carried out as previously described (Brunet *et al*, 2020a). The primary antibodies were diluted as follows: anti-Flag (Sigma, F7425) 1/1000 and anti-GFP (Santa Cruz, sc-9996) 1/8000. The secondary antibodies were diluted as follows: anti-mouse HRP (Santa Cruz sc-516102) 1/10000 and anti-rabbit HRP (Cell signaling 7074S) 1/10000.

### Affinity Purification Mass Spectrometry (AP-MS)

For interactome analysis by mass spectrometry, HEK293 cells at a 70 % confluence were transfected with GFP-tagged PHB or with FLAG-tagged PHBP19 (IP_762813). 24h after transfection, cells were rinsed twice with PBS, and lysed in the AP lysis buffer (150 mM NaCl, 50 mM Tris-HCl and 1 % Triton). Protein concentration was evaluated with a BCA dosage and 1 mg of total protein was incubated at 4 °C for 4 hours with agarose GFP beads (ChromoTek, Germany) for PHB-GFP or with magnetic FLAG beads (Sigma, M8823) for IP_762813-FLAG. The beads were pre-conditioned with the AP lysis buffer. The beads were then washed twice with 1 mL of AP lysis buffer, and 5 times with 5 mL of 20 mM NH4HCO3 (ABC). Proteins were eluted and reduced from the beads using 10 mM DTT (15 mins at 55 °C), and then treated with 20 mM IAA (1 hour at room temperature in the dark). Proteins were digested overnight by adding 1 μg of trypsin (Promega, Madison, Wisconsin) in 100 μL ABC at 37 °C overnight. Digestion was quenched using 1 % formic acid and the supernatant was collected. Beads were washed once with acetonitrile/water/formic acid (1/1/0.01 v/v) and pooled with supernatant. Peptides were dried with a speedvac, desalted using a C18 Zip-Tip (Millipore Sigma, Etobicoke, Ontario, Canada) and resuspended into 30 μl of 1 % formic acid in water prior to mass spectrometry analysis.

### Mass spectrometry analysis of in-house affinity purifications

Peptides were separated in a PepMap C18 nano column (75 μm × 50 cm, Thermo Fisher Scientific). The setup used a 0–35 % gradient (0–215 min) of 90 % acetonitrile, 0.1 % formic acid at a flow rate of 200 nL/min followed by acetonitrile wash and column re-equilibration for a total gradient duration of 4 h with a RSLC Ultimate 3000 (Thermo Fisher Scientific, Dionex). Peptides were sprayed using an EASYSpray source (Thermo Fisher Scientific) at 2 kV coupled to a quadrupole-Orbitrap (QExactive, Thermo Fisher Scientific) mass spectrometer. Full-MS spectra within a m/z 350–1600 mass range at 70,000 resolution were acquired with an automatic gain control (AGC) target of 1e6 and a maximum accumulation time (maximum IT) of 20 ms. Fragmentation (MS/MS) of the top ten ions detected in the Full-MS scan at 17,500 resolution, AGC target of 5e5, a maximum IT of 60 ms with a fixed first mass of 50 within a 3 m/z isolation window at a normalized collision energy (NCE) of 25. Dynamic exclusion was set to 40 s. Mass spectrometry RAW files were searched with the Andromeda search engine implemented in MaxQuant 1.6.9.0. The digestion mode was set at Trypsin/P with a maximum of two missed cleavages per peptides. Oxidation of methionine and acetylation of N-terminal were set as variable modifications, and carbamidomethylation of cysteine was set as fixed modification. Precursor and fragment tolerances were set at 4.5 and 20 ppm respectively. Files were searched using a target-decoy approach against UniprotKB (Homo sapiens, SwissProt, 2020-10 release) with the addition of IP_762813 sequence for a total of 20360 entries. The false discovery rate (FDR) was set at 1 % for peptide-spectrum-match, peptide and protein levels. Only proteins identified with at least two unique peptides were kept for downstream analyses.

### Highly confident interacting proteins (HCIPs) scoring of in-house affinity purifications

Protein interactions were scored using the SAINT algorithm. For each AP-MS, experimental controls were used: GFP alone transfected cells for PHB-GFP AP and mock transfected cells for IP_762813-2F AP. For the PHB-GFP AP, controls from the Crapome repository (Mellacheruvu *et al*, 2013) corresponding to transient GFP-tag expression in HEK293 cells, pulled using camel agarose beads were used. These controls are: CC42, CC44, CC45, CC46, CC47, and CC48. For the IP_762813-FLAG AP, controls from the Crapome repository (Choi et al, 2011) corresponding to transient FLAG-tag expression in HEK293 cells, pulled using M2-magnetic beads were used. These controls are: CC55, CC56, CC57, CC58, CC59, CC60 and CC61. The fold-change over the experimental controls (FC_A), over the Crapome controls (FC_B) and the SAINT probability scores were calculated as follows. The FC_A was evaluated using the geometric mean of replicates and a stringent background estimation. The FC_B was evaluated using the geometric mean of replicates and a stringent background estimation. The SAINT score was calculated using SAINTexpress, using experimental controls and default parameters. Proteins with a SAINT score above 0.8, a FC_A and a FC_B above 1,5 were considered HCIPs.

### Network visualisation of in-house affinity purifications

The network was built using Python scripts (version 3.7.3) and the Networkx package (version 2.4). The interactions from the STRING database were retrieved from their protein links downloadable file. Only interactions with a combined score above 750 were kept.

## Data Availability

The datasets and computer code produced in this study are available in the following databases:

● Protein interaction AP-MS data for both IP_762813 and PHB1 in HEK293 cells were deposited to the ProteomeXchange Consortium via the PRIDE (Perez-Riverol *et al*, 2016) partner repository with the dataset identifier PXD022491.
● Jupyter notebooks containing the analyses are available in the GitHub repository created for this project (https://github.com/Seb-Leb/altProts_in_communities).

## Acknowledgements

We thank the Gygi lab for providing mass spectrometry (MS) datasets and particularly Ed Huttlin for helpful email exchanges. XR, MSS and AAC are members of the Fonds de Recherche du Québec Santé (FRQS)-supported Centre de Recherche du Centre Hospitalier Universitaire de Sherbrooke. This research was supported by CIHR grants MOP-137056 and MOP-136962, and by a Canada Research Chair in Functional Proteomics and Discovery of Novel Proteins to X.R. We thank the team at Calcul Québec and Compute Canada for their support with the use of the supercomputer mp2 from Université de Sherbrooke. We thank Darel Hunting for critically reviewing the manuscript.

## Author contributions

Conceptualization: XR, SL and MAB. Experiments in Fig 1-5, EV1, EV2, data visualization, all Tables: SL. Naive Bayes classifier and interaction scoring: AAC, MSS, SL. Experiments in Fig 6: AML, AD, AT, ABG, MAB and JFJ. Experiments in Fig EV3: MAB and JFJ. Writing_original draft: XR and SL. Writing_review&editing: AAC, JFJ, MAB, MSS, SL, SS. Resources, funding acquisition, project administration: XR. SS and MB initiated the project and mentored SL.

## Extended View Tables Footnotes

*Table extended view 1 - Transcripts and detected altProts for which at least one peptide spectrum match was validated via PepQuery.*

^1^Transcript accessions in bold indicate the longest transcript (used downstream for refProt relative localization).

^2^Biotype that should be assigned given the evidence from the current re-analysis.

^3^If multiple ORFs are present on the transcript and overlap, the transcript is dual coding; if they are sequential the transcript is called bicistronic.

^4^Colored rows indicate pseudogene transcripts that are assigned a multi-coding type.

*Table extended view 2 - Bait-prey pairs involving detected altProts*

^1^A score of 1 indicates that the bait-prey pair constitutes an altProt interacting with the refProt of the same gene, with a shortest path lenght of 1.

^2^A score of 1 indicates that the bait-prey pair constitutes a pseudogene-encoded altProt interacting with the refProt of the corresponding parent gene, with a shortest path lenght of 1. ^3^Set of non-nested (2 aa margin) peptides uniquely mapping to the corresponding altProt.

*Table extended view 3 – CORUM complexes*

^1^Fraction of subunits recovered in the complex.

*Table extended view 4 – altProts coded by pseudogenes for which corresponding parent genes are annotated in psiCUBE (see Materials and Methods)*

^1^ No path indicates that (1) for the pseudogene-encoded altProt, the parent gene-encoded refProt was not identified; or (2) that the altProt and the refProt are not part of the same component in the network.

## Expanded View Figure legends

**Expanded View 1.**
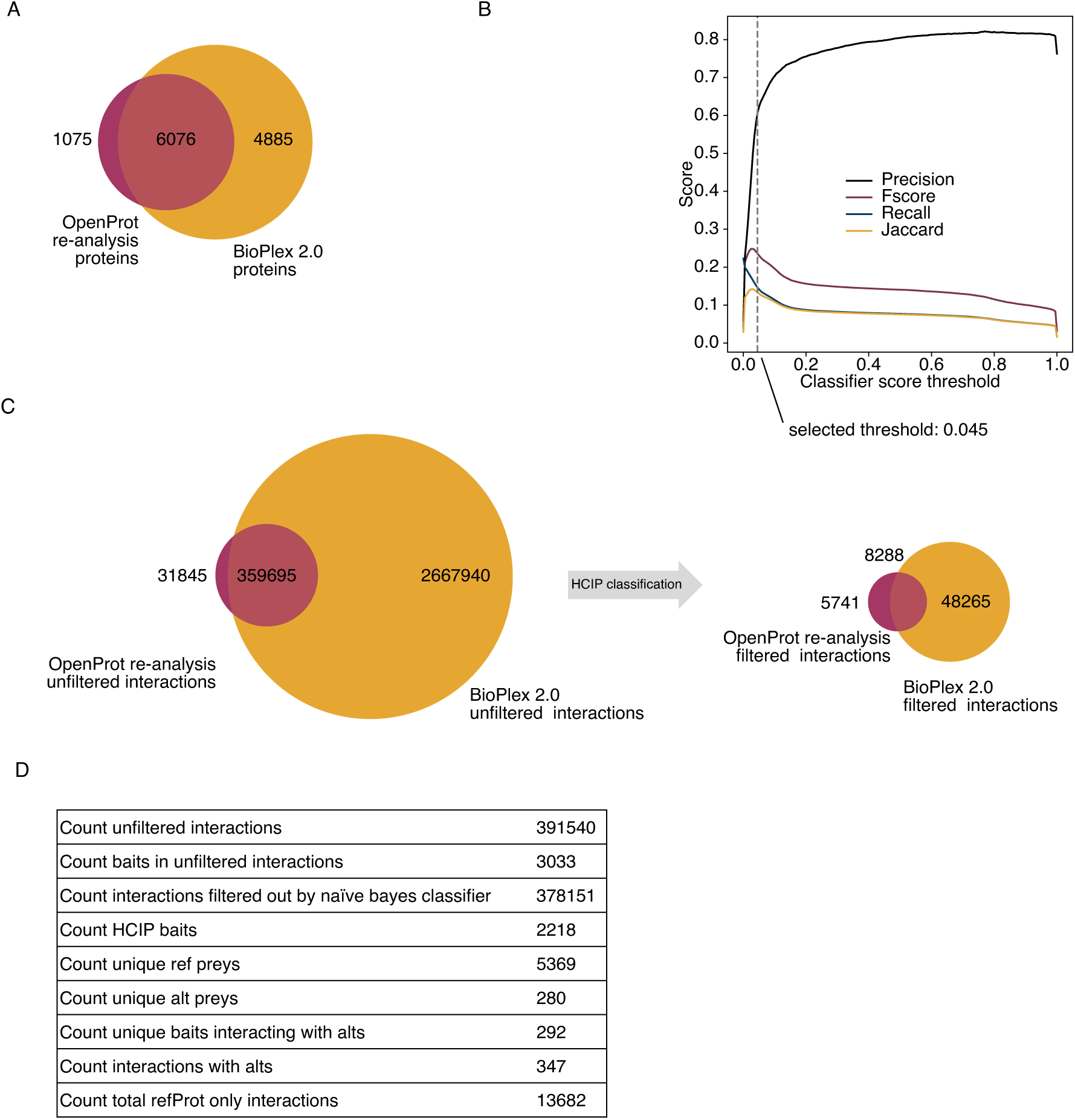
***Network assembly details*** **A** Overlap of total proteins (nodes) in BioPlex 2.0 and OpenProt derived networks. **B** Classifier performance across thresholds. Scores were computed using the BioPlex 2.0 network as ground truth. **C** The overlap of unfiltered interactions between BioPlex 2.0 and the result of OpenProt 1.6 derived re-analysis was considerable (92 % of re-analysis candidate PPIs) (i). Upon filtration the overlap is still significant despite the marked smaller size of the OpenProt derived network (59 % of re-analysis PPIs). **D** Detailed counts of protein and interaction identifications.

**Expanded View 2.**
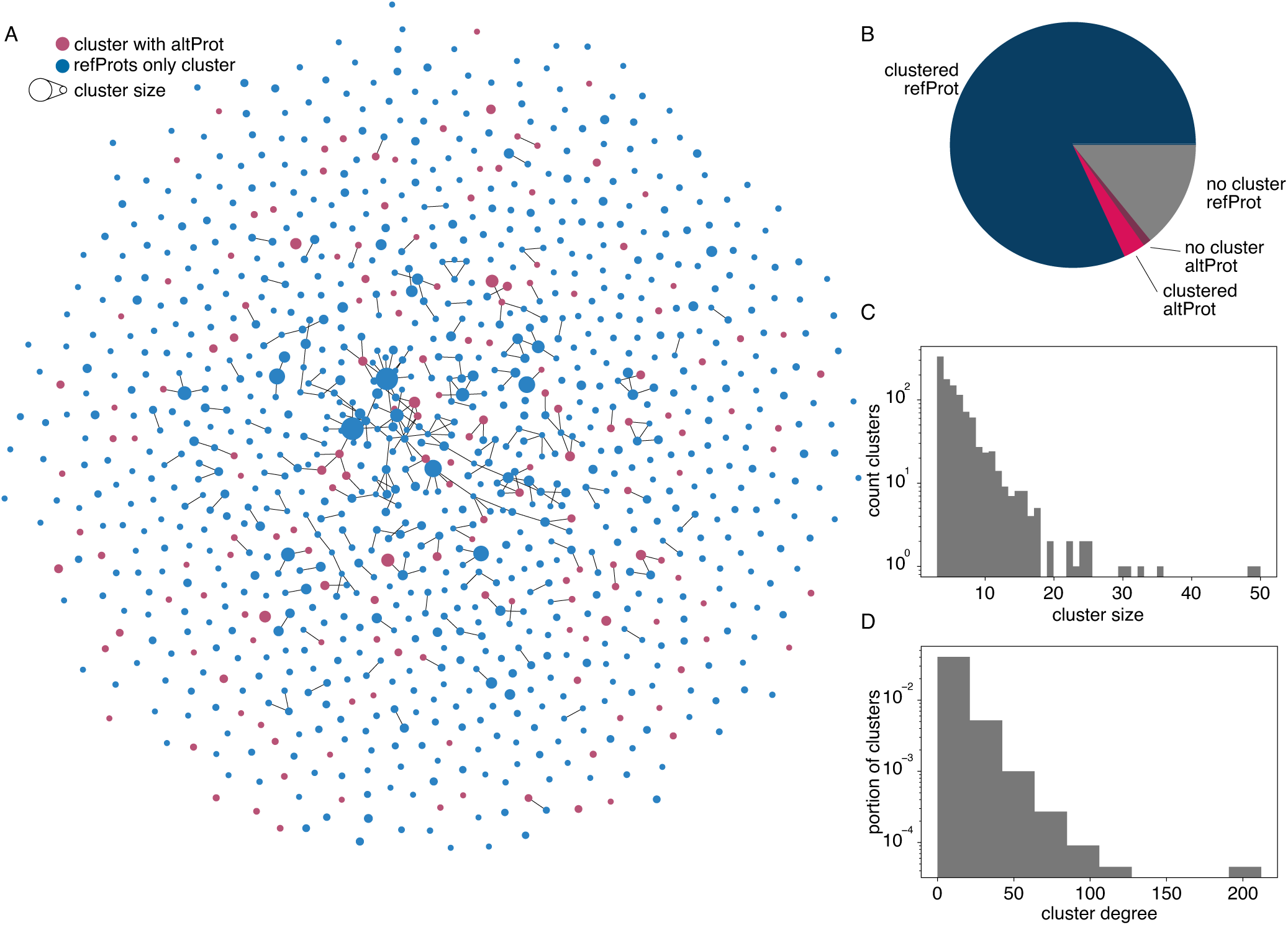
***Community detection details*** **A** Full network of protein clusters. Connections between clusters are drawn if the count of links between their constituent proteins is deemed enriched via a hypergeometric test with alpha set to 0.01 and multiple testing correction set at maximum 1 % FDR. **B** All proteins in the network were either part of a cluster or not and either an altProt or a refProt. **C** Distribution of cluster sizes (count of proteins in clusters). **D** Distribution of cluster connectivity (cluster degree i.e. number of connections a cluster has with other clusters).

**Expanded View 3.**
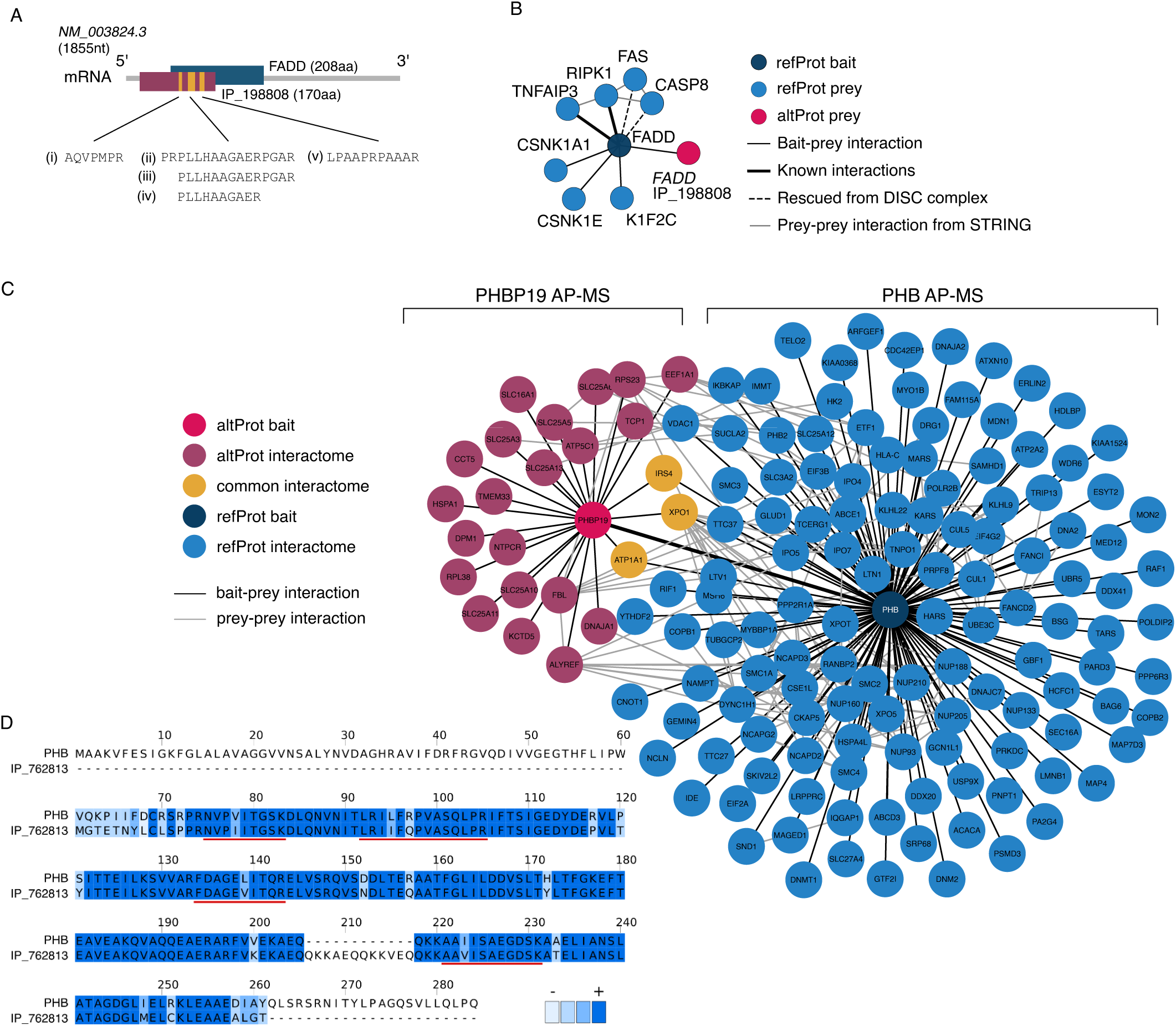
***Validation details*** **A** Validation of interaction between proteins FADD and IP_198808 encoded by the same mRNA. IP_198808 peptides iii, iv, and v were detected in re-analyses of both ViroTrap and BioPlex 2.0 AP-MS of FADD. Peptides i and ii were exclusively identified in ViroTrap and BioPlex 2.0 re-analyses respectively. Peptides spectra matches (PSMs) for i and v from the ViroTrap dataset were validated against unrestricted modifications of reference proteins using PepQuery. **B** FADD network after re-analysis of ViroTrap mass spectrometry data including IP_198808 sequence in the database. **C** Detailed view of the combined network from AP-MS experiments of PHB refProt and PHBP19 altProt. **D** Alignment of IP_762813 altProt encoded by pseudogene PHBP19 and PHB1 refProt sequences based on amino acids using Clustalω with default settings. Blue shading indicates amino acid similarity. Unique peptides detected are underlined red.

## RESPONSE TO REVIEWS

The reviewers provided very constructive comments which we address one by one in detail below. This helped us improve the manuscript. In addition, some comments require additional analyses that we have already performed or that we will perform in a revised manuscript. We are confident that a revised version will satisfactorily address all the central issues.

### REVIEWER #1

In the manuscript “Newfound coding potential of transcripts unveils missing members of human protein communities” Leblanc et al describe an approach to identify and functionally characterise small proteins translated from putative non-coding regions of the human genome (“altProts”). They do this by searching the proteomics raw data of the BioPlex project, which is a large-scale AP-MS project, with candidate altORF sequences and then reconstruct BioPlex’s interaction networks with the newly integrated altProts.

**Major Comment** - I have one major concern about the study, but I appreciate this may be a question for the whole field rather than a single paper. Essentially, I think the most problematic aspect with this kind of experiment is the handling of the false positive detection of small proteins (i.e. the reliable identification of altProts and their interactions), and this appears to be an issue that has not been properly solved yet. The authors clearly err on the side of caution here with stringent cut-offs (0.001% FDR) and additional quality control measures such as testing whether modified peptides of known proteins could have accounted for the newly matched mass spectra. However, I think the authors could describe in more detail how the FDR was calculated, what is the reasoning to bring it down to 0.001% (they only seem to detect 0.1% of all altProt sequences that they search for - so whether this is actually as stringent as it sounds is not so clear given that the search space is huge). Is there a theoretical basis to this and could you elaborate on this in the paper?

***Response*** *- Indeed, this is an important question in the field, and we have explained in previous manuscripts why we currently use such stringent FDR in our re-analyses (e.g. PMID: 32780568, 31033953, 30299502). We’ve added two sentences under Materials & Methods, paragraph Reanalysis of AP-MS data: “**In addition to already annotated proteins, the OpenProt database includes all predicted altProts and novel isoforms. Since large databases result in a large increase of false positive rates (PMID: 23176207, 25357241), this effect is balanced using an FDR of 0.001% as previously described (PMID: 32780568, 31033953)**”*.

*In more details: Large databases are a real challenge in proteogenomics and metaproteomics studies, and several approaches have been proposed, mostly seeking to reduce the size of the database. The most commonly used approach is a two-step database searching method: MS/MS are searched against the large database and PSMs passing a very low stringency scoring (equivalent to a PSM FDR of 0.01%) are used to infer a smaller database. The whole MS/MS are then searched again with this smaller enriched database with a global FDR of 1%*.

*In a recent article, Kumar et al, JPR, 2020 estimated, using entrapment databases, the rate of false positive PSMs at a global 1% FDR for a traditional approach and a two-step approach. The traditional approach yielded a PSM FPR (false positive rate) of 1% in adequation with the global FDR set to 1% but considerably decreased the number of identified proteins. The two-step approach yielded a PSM FPR of up to 10-15% with a global FDR set to 1%*.

*By using a stringent global FDR (0.001%) with a traditional approach, we ensure the rate of PSM FPR is well below 0.01%. By downscaling the global FDR, and thus the PSM FPR, we limit the occurrence of close-but-less-than-perfect PSMs. As a consequence, we obtain a more homogeneous group of PSMs, which allow us to reach a reasonable number of protein identification with high confidence (we limit the noise for the protein inference algorithm). This approach was initially validated by comparison with original studies and manual validations of PSMs (PMID: 30299502). By using this approach, we can be confident of the PSMs, peptides and proteins that are called, and we can also be confident that many are left behind with such a stringent approach*.

**Minor comment 1** - What prior evidence exists in OpenProt that these altORFs are genuine protein-coding elements, e.g. do you shortlist based on ribosome profiling data evidence or such?

***Response*** *- OpenProt displays all predicted altORFs and evidence found to support their protein coding properties including mass spectrometry, ribo-seq, conservation and domain prediction (PMID: 33179748, 302995020). Not all altORFs have such evidence*.

*In this article, we did not shortlist altORFs based on experimental evidence in OpenProt for the following reasons: (1) an absence of detection in OpenProt does not constitute an evidence for absence of expression. (2) BioPlex over-expresses bait proteins which may result in (over) activation of cellular pathways. Thus low abundance protein may be more easily detected in these settings. (3) We ensure confidence in protein identification using combined spectrum- and peptide-centric approaches*.

**Minor comment 2** - Legend 1 F,G section is missing

***Response*** *- Legend 1 F, G section was indeed missing below the figure, but was present in the manuscript file. We are sorry this section was missing in the file provided to the reviewer. This has been corrected*.

**Minor comment 3** - In Fig 2B, only 59.1% of interactions were previously detected by BioPLex. This seems like a lot of new ones (5387 interaction to be precise). Considering these are actually the same raw data, how is that possible and does that suggest that there is a problem with one of the two data processing pipelines?

**Response** - The two pipelines take a different approach to the analysis of the raw data with two major differences that account for the variability:

- ● Difference in protein libraries: BioPlex uses UniprotKB (SwissProt + Trembl), leaving out many reference protein sequences annotated by RefSeq and Ensembl. OpenProt includes these along with all alternative proteins predicted. In total, 35341 protein sequences present in RefSeq or Ensembl but not in Uniprot had a peptide that could be matched to a spectrum found in the raw data.
- ● Difference in peptide-spectrum match (PSM) assignation/ PSM counting and protein inference: BioPlex uses the Sequest search engine to find peptide spectrum matches considering fully tryptic peptides that are then assembled into proteins using their in-house protein inference tool. The OpenProt pipeline uses four search engines via SearchGUI, including search algorithms that consider non fully-tryptic peptides, and PeptideShaker for protein inference from peptide identification.

These differences account for the different PSM counts used to train the classifier and thus resulted in a different overall network.

It does raise the problem of variability of results depending on the method of analysis starting from the same raw data, as previously observed (PMID: 33133425), but there doesn’t seem to be a consensusof superiority either way.

**Minor comment 4** - Fig 2E: The eigenvector centrality measure is an interesting idea, but doesn’t the plot suggest that altProts are enriched towards the left, i.e. lower EVC? I think I’m missing something here in the description of the figure…

***Response*** *- The reviewer is correct: altProts seem enriched towards the left, i.e. lower EVC. However, while altProts are more often seen in the periphery, a significant fraction is found in central regions of the network. In addition, since no altProts were used as baits in BioPlex 2.0, they are likely artificially pushed towards the edges of the network. We have added the following sentence, page 11: “**Since no altProts were used as baits, they are likely artificially pushed towards the edges of the network**”*.

**Minor comment 5** - It’s quite striking that BioPlex3 dismissed 20% of their previously claimed interactions in BioPlex2. That sounds like the protein-protein link FDR is unacceptably high in BioPlex. Do the authors think that their more stringent setup is better in this respect and could they show evidence to support that?

***Response*** *- Our approach is indeed more stringent at the protein identification step. This setup was necessary for confident identification of novel proteins. The stringent FDR is to compensate for the larger protein library used to search spectral matches (see response to comment 1 above). As expected, this strategy results in a different profile of PSM counts overall (i.e. total number of PSMs, identity), compared to BioPlex 2.0 who affects the training of the classifier that identifies HCIPs. However, we expect that the FDR at the PPI level is comparable to the BioPlex 2.0 since the interaction identification pipeline downstream identification was closely replicated*.

*We suspect that the difference between BioPlex 3.0 and 2.0 also comes from a different profile of PSM counts as the authors re-ran all their MS/MS data analysis in BioPlex 3.0*.

**Minor comment 6** - Please explain the term “protein communities” earlier on in the paper

*Response - We have added the following sentence in the third paragraph of the Introduction: “Here, a community represents a group of nodes in the network that are more closely associated with themselves than with any other nodes in the network as identified with an unsupervised clustering algorithm”*.

**Minor comment 7** - GAPDH: how different are the detected peptides of the pseudogenes and how many peptides are there for each? In other words, how sure can you be that these are actually the pseudogenes you detect, and not just affinity-purified canonical GAPDH.

***Response*** *- All peptides identifying each pseudogene are unique: they only identify a pseudogene and not the canonical GAPDH nor another pseudogene. Furthermore, all peptides uniquely map to each pseudogene and have been confirmed with PepQuery. This peptide-centric algorithm verifies that the experimental spectra is not better explained by a known protein with any post-translational modification. We will include some of the spectra in a revised manuscript*.

**Minor comment 8** - The abstract is not so clear about the methodology. It should specify that these networks are protein-protein interaction networks in my opinion.

***Response*** *- We agree, and we have made two modifications:*

- *We changed “communities” to “**protein-protein interaction networks**” in the second sentence*.
- *We also changed the third sentence “Here we incorporate this increased diversity in the re-analysis of a high throughput human network proteomics dataset thereby revealing the presence of 203 alternative proteins within 163 distinct communities associated with a wide variety of cellular functions and pathologies” to “**Here we used the proteogenomic resource OpenProt and a combined spectrum- and peptide-centric analysis for the re-analysis of a high throughput human network proteomics dataset thereby revealing the presence of 280 alternative proteins in the network**”*.

**Minor comment 9** - I think you should mention the work of the sORFs.org team as well

***Response*** *- We have added the following reference (lane 87): Olexiouk V, Van Criekinge W, Menschaert G (2018) An update on sORFs.org: a repository of small ORFs identified by ribosome profiling. Nucleic Acids Res. 46(D1):D497-D502*

**Minor comment 10** - Page 7, line 149: “RefProts from 4656 genes (or 86% of total re-analysis results) were found in both the BioPlex 2.0 and in the present work (Fig EV1A), indicating that the re-analysis could reliably reproduce BioPlex results.” The figure shows different numbers. Also, 86% overlap in identifications is low - does that mean that the 1% FDR for protein identifications in BioPlex was actually more like 14%?

***Response*** *- The reviewer is right to point out the rounding error, we have changed the number to **85%** in the manuscript*.

*Considering the differences in the analytical pipeline, 85% overlap is actually quite reasonable (PMID: 30299502, 33133425)). This figure does not mean that BioPlex obtained a higher FDR, it means that in the BioPlex pipeline some spectra were left unassigned because the matching peptides were not in the library. This accounts for the identifications present in the re-analysis but absent in the original study. For the ones present in the original study but absent in the re-analysis the discrepancy can be explained by the extra stringent FDR of 0.001%*.

Reviewer #1 (Significance (Required)):

**Other comment** - I think this is an interesting manuscript and approach, and very timely. To my knowledge, many groups are currently interested in detecting translation from “non-coding” genomic regions, but few methods exist that enable the functional characterisation of such proteins. To me this is possibly the key aspect the paper and could be fleshed out more. For example, the communities and their GO enrichments given in Figure 4 could be presented additionally in a format that is more easily accessible as a resource, e.g. a supplementary table or simple website.

***Response*** *- We thank the reviewer for this comment and the suggestion to present some data in a more easily accessible way. We had already planned to add a webpage to the OpenProt proteogenomic resource that displays protein communities and GO enrichments:* http://openprot.org/ppi*. This webpage is under construction and will be up and running within the next 4 weeks*.

My expertise is in proteomics data analysis.

### REVIEWER #2

**Summary:**

The manuscript “Newfound coding potential of transcripts unveils missing members of human protein communities” submitted by Leblanc et al reports on a reanalysis of the BioPlex AP-MS dataset with the aim to detect evidence for novel proteins and their interactions with known proteins. Reference protein databases used to match MS spectra to peptides and proteins usually only consist of the canonical proteins and well-described isoforms. Thus, peptides from alternative ORFs would not be detected. Leblanc et al performed a search against a much larger protein reference database called OpenProt and indeed detected a few hundred alternative proteins and their interactions with known proteins. The authors subsequently analyze their data primarily with respect to the putative interaction partners of these altProts performing various network analyses and finally experimentally validating selected altProt-knownProt interactions using coIP and imaging in over-expression systems. The authors conclude that translation of alternative ORFs in human is widespread and likely results in the production of functional proteins as they can interact with other known proteins. They suggest that their data can be the starting point for experimental investigations into the functions of these altProts and that human genome annotation should allow for polycistronic gene models. Because a fraction of the identified altProts (i.e. from pseudogenes) interact with their refProt, they further suggest that one way for how altProts might function is to lead to altered functionalities of refProts that can form homodimers or heterodimers with paralogs but also with proteins from pseudogenes. The manuscript is very clearly written and the figures are overall very nice.

*We thank the reviewer for this thorough and fair summary*.

**Major comment 1** - The conclusions drawn in this manuscript seem accurate based on the data presented with two exceptions.

1. Without being an expert in gene expression and genetics, it remains unclear how the authors can differentiate whether two ORFs from the same gene that they either classified as dual gene or bicistronic actually rather represent a discovery of new alternative isoforms of a gene. To the best of my knowledge, alternative isoforms can be partially overlapping or non-overlapping at all questioning the request of the authors in the discussion to open up the human genome annotation to polycistronic genes. The manuscript would benefit from a more detailed description for how and why the authors came up with their classification of altProts and the corresponding genes.

***Response*** *- Our annotations use transcript sequences as the starting point. From there all open reading frames of 30 codons or longer are predicted as potential coding sequences. Protein isoforms derived from alternative splicing typically share some sequence similarity with the canonical protein with alteration based on the configuration of exon and intron excision/retention. For these isoforms, we use BLAST search filtered fora bit score over 40 for an overlap over 50% of the queried reference sequence (PMID: 33179748). We can confidently identify altProts as novel protein products because they share no sequence similarity with canonical proteins of the same gene*.

*We have added a paragraph in the methods section (page 26) to clarify the classification of proteins, transcripts and genes:*

*Classification of proteins, transcripts and genes*

*Reference proteins (RefProts) are known proteins annotated in NCBI RefSeq, Ensembl and/or UniProt. Novel isoforms are unannotated proteins with a significant sequence identity to a RefProt from the same gene. These isoforms are identified with a BLAST search filtered for a bit score over 40 for an overlap over 50% of the queried reference sequence. Alternative proteins (AltProts) are unannotated proteins with no significant identity to a RefProt from the same gene. Importantly, altProts may share a sequence similarity with a protein from a different gene, for example in the case of pseudogene-encoded altProts and the protein derived from the parental gene*.

*Alternative open reading frames (altORFs) correspond to unannotated ORFs predicted to encode proteins with no significant identity to any other annotated protein*.

*We classify RNA transcripts as dual coding or bi-cistronic based on the relative position of the ORFs on the transcript. If they are overlapping (i.e. if they share nucleotides) we classify the transcript as dual coding, if they are sequential (i.e. share no nucleotides) we classify it as bicistronic. Gene classification with this respect is inherited from the classification of transcript it produces. Note that transcripts and genes can hold both dual coding and bi-cistronic classifications*.

**Major comment 2** - Line 512-515: The authors state that the impact of altProts on network structure is revealed by the bridging role many seem to play… There is no proof for this statement. To show this, the authors would need to go beyond a visual inspection of the network data and perform computations such as removal of altProts from the network results in more disconnected components than random removal of refProts (degree-controlled, of course). Also, the example in figure 3Ai does not show that the complexes are otherwise independent/not connected.

***Response*** *- The claim is not that altProts play the role of bridges more often than do refProts (which is what the reviewer suggests to test). Only that in some cases altProts bridge otherwise unconnected or sparsely connected regions. This is not an overstated claim, the simple presence of alt nodes with eigenvector centralities higher than most ref nodes indicates the contribution of at least some alts towards overall network connectivity. We maintain that our stringent identification of several altProts in the purification of multiple baits (e.g. IP_117582|BEND4 identified in 7 purifications) does lend support to the claim that network structure is altered (i.e. different nodes & different edges) when considering the presence of alternative proteins. The bridging role observed simply points to the fact that many altProts are the only direct link between two or more baits with which they purified. A simpler example is Figure A which shows that no path shorter than 4 is possible between the two baits in the overall network (since all their direct interactors and the edges between them are present in the subnetwork) but the simple addition of the altProt connects them with a path length of 2*.

**Major comment 3** - The description of the experimental work lacks some detail. I.e. the transfections used are not coupled with the actual experiment, i.e. coIP, meaning that the reader has to infer how cells where transfected for which type of experiment. Also, the transfection section mentions siRNA experiments. Where in this study were siRNA transfections been conducted?

***Response*** *- The Material and Methods is grouped by methods which seems to be in adequation with previous articles published in Molecular System Biology journal. The comment of the reviewer is somewhat unfair as all details particular to specific experiments have been pointed in each of the relevant sections (e.g. “For co-immunoprecipitation of PHB1-GFP and RPL18-GFP, stringent wash were done with modified lysis buffer (250 mM NaCl + 20 μg/ml peptide FLAG (F3290 Sigma)) prior to elution with 200μg/ml peptide FLAG.”). Since the same method (unless otherwise stated) was used for all experimental validations, organizing the Material and Methods by experiments does not seem ideal to us, although we could reorganize it should this be wished by the editor*.

*As far as the comment on siRNA, we did not perform siRNA experiments in the manuscript. Maybe there was a confusion with the name of our transfection reagent, jetPRIME®, DNA and siRNA transfection reagent (see:* https://www.polyplus-transfection.com/products/jetprime/).

**Minor comment 1** - The fact that quite some altProts interacted with their refProt was quite puzzling and interesting. The hypothesis as presented by the authors that the refProts might engage in homodimers and heterodimers with paralogs is logical, however, the manuscript would have benefited from a more thorough analysis in this direction, also because it is one of the key findings of the paper, as it seems. One would assume for example that altProt-refProt interactions primarily occur with altProts from pseudogenes or where altProts when from the same gene as the refProt still share some exons or are partially in the same frame. Is there any trend in this direction?

***Response*** *- We agree with the reviewer that it is a particularly puzzling observation, in particular since it does not seem to correlate with the degree of protein sequence identity for pseudogenes (figure 2D). Exons are indeed shared (i.e. between FADD and altFADD) but the respective amino acid sequences are completely different because they are translated from different reading frames. While some of the interacting pairs seem to indicate that sequence identity is the driving factor (i.e. pseudogene-parent genes), others indicate other modes of interactions are at play. It is also interesting to note that refProt-altProts duos share a transcriptional and post-transcriptional regulation since they are present on the same transcript while it is not the case for the parental gene-pseudogene couples*.

**Minor comment 2** - To better understand the identified altProt-refProt interactions, it would have been helpful for the presented candidates to systematically indicate what the sequence similarity was to rationalize whether a heterodimer kind of mechanism is plausible or not. In the same line, highlighting which altProt-refProt candidates in the figures are unlikely to occur based on a heterodimer mechanism would have also been very interesting. My impression is that this information is difficult to retrieve from the current data provided.

***Response*** *- We assume this question is about pseudogenes-derived altProts as they may display a high degree of similarities with proteins coded by the corresponding parental genes. This comment also relates to reviewer #3 comment 1: “It would have been better if the authors had shown the amino-acid alignments of the ref and altProts identified here in the Supplementary Data. It is impossible to follow how much the ref and corresponding altProts differ in respect of their sequence”*.

*We will provide these alignments data in a revised manuscript*.

**Minor comment 3** - The authors refer to altProts that are encoded by pseudogenes. GENCODE classifies pseudogenes in a variety of different classes, i.e. not-transcribed, transcribed, translated, polymorphic, processed, etc. For a better understanding of the origin of the identified altProts, it would have been helpful to further analyze whether they tend to originate from a specific subclass of pseudogenes.

***Response*** *- OpenProt annotations start from the annotated transcriptome (Ensembl and RefSeq). Hence, pseudogenes-derived altProts are obligatorily predicted from transcribed pseudogenes. As suggested by the reviewer, we will provide a detailed classification of pseudogene classes identified in a revised manuscript*.

**Minor comment 4** - I wonder whether more orthogonal data could have been used to further annotate and substantiate the identified altProts. This would also increase the value of the data as a resource to prioritize altProts for further experimental validation. Would it be possible to search in tissue transcriptome datasets like GTEx for example for further transcript evidence of the altProts or in alternative proteome datasets like Wilhelm et al Nature 2014 or Kim et al Nature 2014? Is there more evidence from external sources for altProts that the authors identified in their study compared to randomly selected other altProts from OpenProt?

***Response*** *- 156 altProts in the network showed additional MS evidence from datasets other than BioPlex on the OpenProt online resource. 18 altORFs encoding altProts in the network show evidence of translation initiation/elongation via ribo-seq on OpenProt online ressource. We will include this information as additional columns to Table EV2 in a revised manuscript*.

*As far as searching in tissue transcriptome datasets like GTEx, OpenProt starts from transcriptome assembly (Ensemble & RefSeq), hence all altProts have evidence at the transcript level (= at least one transcript with experimental evidence contains the coding sequence in question)*.

**Minor comment 5** - Fig 2c: I don’t think the data shows a power law because the data does not show a linear correlation, which is something that I have observed before for BioPlex and other AP-MS data probably because hubs are filtered out as non-specific binders leading to a kind of plateauing.

***Response*** *- Visually the distribution seems to diverge from a power law at the extremities but the maximum likelihood method indicates that it actually tends toward that distribution. There seems to be a lack of both small degree and high degree nodes. As the reviewer mentions, a lack of higher degree nodes induces a plateau like shape towards the right, but this is likely due to the asymmetrical nature of AP MS data where true hubs are better discovered if they are used as baits. Conversely, the lower degree nodes are lacking because the stringent filtration likely erodes a number of true positives from the unfiltered dataset*.

*In an ideal set-up all proteins would be used as baits to have a complete network that then would fit a power law distribution. Between, the absence of some important baits (hubs), a stringent filtration to avoid false positives and an incomplete experimental network (not all proteins were baits), it is not surprising at all that the distribution shall diverge from a power law distribution, in particular at the extremities*.

**Minor comment 6** - Line 251: What do the authors mean with “neighborhood”? Please, specify.

***Response*** *- We mean “directly interacting with one or more subunits in the complex”. The sentence “We observed 50 altProts in the neighborhood of CORUM complex subunits that served as bait” was changed to “**We observed 50 altProts in the neighborhood of CORUM complex subunits that served as bait, i.e. directly interacting with the CORUM complex**”*.

**Minor comment 7** - Line 273: Typo: Theubiquitin

***Response*** *- Thank you; this was corrected*.

**Minor comment 8** - Line 287-290: This statement seems out of context and it is unclear what the link of the data is to tumorigenesis.

***Response*** *- Our data only brings forth the fact that ELP6 has an unannotated interacting partner that may lead to a better understanding of its function. Although no direct link to tumorigenesis can be made from our analysis, our claim is simply that IP_688853 should be part of further investigations surrounding the involvement of ELP6 in the pathological process*.

**Minor comment 9** - Line 319-320: The analyses of the sequence similarities are somewhat unsatisfactory without stating what a reasonable cutoff of sequence similarity should be for example. It doesn’t require high sequence similarity to maintain the same fold and still be able to heterodimerize for example. Figure 3Di is not very helpful because it is hard to interpret the alignment score. Why not using a more simple quantification like fraction of identical residues with respect to the length of the altProt sequence?

***Response*** *- Indeed it is not necessary to have high sequence similarity to maintain fold and heterodimerize, as Fig 3Di (now Fig 3D) shows with pairs of proteins directly interacting are present throughout the range of Needleman Wunsch (NW) alignment score. NW score is computed similarly to what the review suggested: it is a local assessment of sequence similarity with penalty on gaps and mismatches*.

**Minor comment 10** - The numbering of figures appears unusual at times. Authors can consider changing some i numberings to an actual capital letter, i.e. Dii to E.

***Response*** *- We agree with the reviewer and we have changed Dii to E. As for the other figures, we believe that the current numbering is OK*.

**Minor comment 11** - Line 324-333: This paragraph could be shortened to one or two sentences.

***Response*** *- This paragraph is already pretty dense and the provided information all relevant. We could not find a way to shorten this paragraph without omitting important information*.

**Minor comment 12** - Fig 4A. The hypergeometric test might not be appropriate here but it is difficult to assess with the information provided in the methods. Did the authors take into account the different degrees of proteins in different clusters? Usually, significances of connectivity between two groups of genes is assessed empirically using degree-controlled randomized networks.

***Response*** *- The hypergeometric test is used here to assess the significance of an enrichment. In this case it is the set of edges connecting nodes . In this respect the node degree is taken into account. The exact methodology was reproduced from Hutlin et al 2015*.

**Minor comment 13** - Fig 4B. Using all human genes as background for the GO enrichment analysis might not be appropriate. Wouldn’t it make more sense to use all proteins in the dataset as background to avoid enrichments just because some proteins are more amenable to AP-MS than others?

***Response*** *- Our rationale behind the use of the whole genome is that BioPlex is a high throughput survey of interactions in the whole proteome. Several methods have been used in the literature for computing gene set enrichment statistics with different backgrounds including whole genome, input set and filtered set. As suggested by the reviewer, we will add the enrichment analysis considering all identified proteins as background to a revised manuscript with possible alterations to figures 4 and 5*.

**Minor comment 14** - Line 437. Please, provide some detail in the results section on how cells were transfected (i.e. with which fusion constructs).

***Response*** *- We have provided these details as suggested by the reviewer*.

- *Page 19: we changed “In cells, FADD formed larged filaments…” to “In cells co-transfected with Flag-FADD and IP_198808-GFP, FADD formed large filaments…”*.
- *Page 20: we changed “The interaction between altProt IP_624363 encoded in the EEF1AP24 pseudogene and EEF1A1 (Fig 3Av) was confirmed by co-immunoprecipitation…” to “**The interaction between altProt IP_624363 encoded in the EEF1AP24 pseudogene and EEF1A1 (******Fig 3Av) was confirmed by co-immunoprecipitation from cell lysates from cells co-transfected with GFP- eEF1A1 and IP_624363****…”*.
- *Page 20: we changed “First, PHB1 co-immunoprecipitated with IP_762813…” to “First, PHB1 co-immunoprecipitated with IP_762813 using cell lysates from cells co-transfected with PHB1-GFP and IP_762813-Flag…”*.
- *Page 20: we changed “The interaction with RPL18 was tested and confirmed by co-immunoprecipitation (Fig 6D, left)…” to “The interaction with RPL18 was tested and confirmed by co-immunoprecipitation in cells co-transfected with RPL18-GFP and IP_117582-Flag (Fig 6D, left)…”*

*Furthermore, we have generated a supplementary table containing all nucleotide and protein sequences of the transfected constructs. This table will be added to the revised version of the manuscript*.

**Minor comment 15** - Fig 6C. Why was there no imaging/co-localization experiment performed as it was done for the other presented candidates? If this is because the experiment did not work, then it is ok to state this and rationalize why you then chose a different experimental approach. The authors should also report how many altProt-refProt interactions they in total assessed and how many of them were validated.

***Response*** *- We did not include these initially as the over-expression of PHB leads to mitophagy and results in a collapsed mitochondrial network around the nucleus. However, these experiments were performed and showed a colocalisation between PHB and PHBP19 (IP_762813). We will add these in the supplementary figure of the revised manuscript as we agree with the reviewer that it is still worth showing*.

**Minor comment 16** - Data availability: Do you think your data would qualify to update human genome annotation? If so, the authors should consider submitting their data to GENCODE for example, if possible, or at least state how many genes, in their opinion, should change their annotation.

***Response*** *- Protein sequences have been submitted to GenBank and a third party annotation accession number will be available shortly*.

Reviewer #2 (Significance (Required)):

**Significance:**

**Other comment 1** - In the minor comments section I suggested a couple of more analyses which in my opinion might significantly increase the value of the manuscript. Currently, apart from the reanalysis of the AP-MS data, the manuscript does not seem to present other major novelties in the field of alternative ORFs in human and the authors don’t specify or provide sufficient information how it can be used to improve human genome annotation or functional characterization of altProts. However, I am also not following in detail the field of de novo protein detection in human and human genome reannotation.

***Response*** *- To the best of our knowledge, this is the first time that altORFs or smORFs have been shown to be extensively present and involved in the human interactome on such a large scale. We believe that this article will represent a landmark in the community where scientists will dig for hypotheses to be tested in the lab, but also as it showcases the role of altProts in human PPIs*.

*We have submitted to GenBank the sequence of the 295 altProts detected in our study (Figure 1C)*.

**Other comment 2**- I cannot judge the accuracy of the MS data reanalysis and whether enough evidence has been presented for the existence of the altProts in the MS dataset.

**Response** - We used a highly stringent spectrum-centric approach (FDR 0.001%) combined with a stringent peptide-centric approach (PepQuery). PepQuery is a peptide-centric algorithm validating that each experimental spectrum is not better explained by any random peptide, unmodified canonical peptide or a canonical peptide with any post-translational modification. We will also provide some MS spectra in response to some comments from reviewer #1 and reviewer #2.

### REVIEWER #3

Reviewer’s comments on the manuscript by Leblanc et al. “Newfound coding potential of transcripts unveils missing members of human protein communities”

First of all, I would like to apologise for the delay in reviewing this manuscript. Leblanc et al. re-investigated mass spectrometric data from large-scale affinity-purifications of human proteins that are available in the BioPlex 2.0 network in order to identify hitherto non-identified protein forms, so-called altProts. For this, the authors used their recently published OpenProt proteogenomics library and the OpenProt MS pipeline in order to identify proteins, including their sequences, which are encoded by alternative open reading frames (altORFs) and lead to translation of altProts. Matches obtained at very stringent FDR settings were validated with PepQuery.

In the Bioplex 2.0 dataset the authors found a number of proteins that are not yet included in reference databases (refProts), encompassing proteins derived from pseudogenes, ncRNAs and alternative ORFs of canonical and ref mRNAs. Furthermore, the authors used their data to rebuild a dataset/network by employing their identified altProts as prey proteins and using the CORUM database to assess the portion of complex subunits in their novel network. Here they found interesting contributions from their identified altProts.

The authors also validated in a functional manner three of the altProts, each derived from a different group: (i) a dual-coding gene (FADD coding gene), (ii) a pseudogene, (iii) a different gene. The authors used co-affinity purifications and immunofluorescence to monitor protein interactions between the alt and refProt as well as the different locations of FADD, EEF1A1, PHB1 and BEND4 and its corresponding altProts.

In general, I very much appreciate the authors’ efforts to expand the (human) proteome by the reliable identification of proteins which have sequences different from those of the current reference proteins present in the databases. However, I have several points that need to be clarified before this work can be regarded as suitable for publication.

***Response*** *- We thank the reviewer for his appreciation of our efforts to identify altProts in the interactome of refProts, an important step towards the determination of their molecular functions. We address the points raised by the reviewer below*.

**Major comment 1** - It would have been better if the authors had shown the amino-acid alignments of the ref and altProts identified here in the Supplementary Data. It is impossible to follow how much the ref and corresponding altProts differ in respect of their sequence.

***Response*** *- Within an mRNA, the annotated CDS and the altORF are two completely different coding sequences. However, we acknowledge that visual amino-acids alignments may be helpful to show that refProt/altProt pairs encoded in the same gene have different sequences. We will provide these alignments data in a revised manuscript. For altProts coded by pseudogenes, the difference between the parental and the pseudogene-derived protein could be as subtle as a change in a single amino acid change to a significant change such as the deletion of an entire domain. Here too, the alignments will be provided in a revised manuscript*.

**Major comment 2** - The underlying database of predicted ORFs has been shown to be suitable in the detection of altProts in single-case studies, which have been validated by cell-biological or biochemical experiments. However, the application of the database in high-throughput studies, like the one presented in this manuscript, has not been shown or validated so far (at least I could not find any example in the literature). Could the authors point to relevant studies where this approach has been benchmarked or validated? In case such study does not exist so far, it would be important to include such validation in the current manuscript. The following points 3-6 are suggestions that I consider as minimum requirement for validation of the MS analysis workflow.

***Response*** *- To follow-up on this comment, we would like to point out that there are previous examples in the literature with application of the OpenProt database in high-throughput studies (e.g. PMID: 32891891, 33352703, 33133425). OpenProt itself re-analyzes large proteomics datasets using its own database to add experimental evidence to alternative proteins from different organisms (PMID: 30299502, 33179748). In addition, the high throughput nature of the study would not affect the ability to identify peptides in mass spectral data using the OpenProt database; in such studies, a variable number of mass spectrometry RAW files are re-analyzed with the OpenProt database. The only difference between the re-analysis of small-scale and large-scale studies would be the computing time*.

**Major comment 3** - The authors aim to encounter the significantly larger peptide search space for altProts compared to canonical proteins by applying a very stringent protein FDR. It is well known that MS database search engines strongly underestimate FDRs when challenged with very large search spaces. Therefore, application only of a very stringent FDR is not sufficient. At least when working with FDRs the authors must apply an FDR filter at the peptide level. This is even more important, since the authors seemed to have identified many peptides derived from altProts that differ only in one or two amino acids to peptides derived from canonical proteins.

***Response*** *- Typically, an altProt encoded in an mRNA has a completely different amino acid sequence from the annotated protein because the altORF is different from the annotated ORF. In contrast, an altProt encoded in a pseudogene-derived ncRNA may be very similar to the parental protein and a specific peptide from this altProt may differ only in one or two amino acids, as indicated by the reviewer*.

*We agree with the reviewer that the use of large databases such as OpenProt calls for cautious approaches. Here, we used a highly stringent spectrum-centric approach (FDR 0.001%) combined with a stringent peptide-centric approach (PepQuery). A more detailed explanation on the need and consequence of the stringent FDR is provided in answer to the first comment of reviewer 1. Of interest, PepQuery is a peptide-centric algorithm validating that each experimental spectra is not better explained by any random peptide, unmodified canonical peptide or a canonical peptide with any post-translational modification*.

**Major comment 4** - The authors consider a database of predicted ORFs merged with the canonical proteome database to assign peptide spectrum matches (PSMs). It is known that large proportion of peptides derived from canonical proteins can be mapped to non-canonical genomic regions (such as the predicted ORFs in the applied database of this study) and vice versa. Due to this sequence similarity, the mapping of peptides to their ’true’ origin poses an enormous challenge. The authors should also consider “alternative mappings”, which are not yet included in the database of predicted ORFs. Those include for example peptide derived from frame shift events or canonical peptides carrying single amino acid mutations. All theoretically alternative peptide sequences could be determined by in silico translation of genomic sequences. Subsequent alignment of the identified peptide sequences derived from altProts against the resulting in silico translated database can be done fast, for example with tools such as ProteoMapper (Mendoza et al, 2018, Journal of Proteome Research).

***Response*** *- In silico translation of an entire genome (3-frame or 6-frame) as suggested by the reviewer would result in millions of protein sequences and billions of peptides. No search engine with any overly stringent FDR would resolve such search space (see response to comment 1 of reviewer 1). That is why OpenProt starts with transcriptome annotations and performs a 3-frame in silico translation. Hence, any “alternative mappings” with a transcript support would be considered in our analysis. Because we don’t start from the genome, but from the transcriptome, we already include frame-shits from all alternative splicing events with evidence at the transcript level. It remains the question of SNPs, but here we work with a cultured cell line that is well characterized. Admittedly when working with biological material from a different origin, it may be desired to sequence the genome or the transcriptome to have a personalized database*.

**Major comment 5**- I appreciate the authors’ re-evaluation using their OpenProt database using stringent settings and validation with PepQuery. However, I definitely consider PROSIT as a viable alternative, as it is not FDR-based, and I recommend that the authors validate the identified peptides of altProt by PROSIT. I suggest the authors predict for both, canonical and altProt derived peptides the MS2 fragmentation pattern using PROSIT, and subsequently compare detected and predicted MS2 for example via dot products between spectra. The resulting distributions of dot products for peptides derived from altProts should assemble the distribution of dot products for peptides derived from canonical proteins. A difference in distribution would point to differences in FDR for canonical vs altProt peptides.

***Response*** *- We appreciate the suggestion of the reviewer, however PROSIT requires a MaxQuant input (msms.txt), when our analysis uses SearchGUI/PeptideShaker to take advantage of multiple search engines. However, we believe PepQuery (PMID: 30610011) offers a similar analysis to what the reviewer suggests : for each queried peptide, a spectra validated by PepQuery indicates that this spectra is better explained by the queried peptide, than by any random peptide, unmodified canonical peptide or a canonical peptide with any post-translational modification*.

*Hence, altProts are identified in a manner even more robust than the refProts: any bias present is against altProt identification*.

**Major comment 6** - The authors performed validation studies of exogenously expressed altProts. To convince me further that indeed these altProts are expressed I would like to have those altPep and their corresponding MS2 spectra identified and hence used for confirmation (under stringent FDR settings) that the authors indeed validated these (or at least some of the) altPep and their MS2 spectra by comparison with those the corresponding synthetic peptides.

***Response*** *- The validation studies were primarily performed to confirm protein-protein interactions and colocalization, not to confirm the expression of altProts*.

*We agree that a confirmation of the endogenous expression of some of these altProts using synthetic peptides is needed, and we are already planning such experiments for some altProts. However, we feel that this is out of scope for the current study*.

**Major comment 7** - I do not understand the goal of the authors do rebuilt a novel network based on available Bioplex 2.0 data? If this was because the network had major shortcomings – e.g., if the absence of altProts led to a completely different protein interaction network – then I would have understood this. However, as far as I can see, the authors used their OpenProt database and re-evaluated the AP-MS data with more stringent settings (mainly mass deviation and FDR). Surprisingly, they built a novel network that overlaps with the Bioplex network to less than 60%. For me, this low degree of overlap implies a completely different network and thus should be another main message of the authors’ manuscript. Therefore, the authors should compare the Bioplex 2.0 or 3.0 networks with their own network more thoroughly, and they should describe the difference in more detail.

***Response*** *- It was necessary to rebuild the network because identification of protein protein interactions in BioPlex relies on PSM counts of the whole study at once to train the classifier that filters out background to identify HCIPs. We could not simply place identified altProts in the already existing BioPlex network because the simple identification in AP-MS does not necessarily imply protein-protein interaction, as stated in other comments of the reviewer*.

*We will include a more in depth discussion on the differences between the OpenProt-derived, BioPlex 2.0 and BioPlex 3.0 networks in a revised manuscript*.

**Major comment 8** - I cannot entirely follow the authors’ argument that an altProt interacts with a refProt and thereby generates a novel network and/or alternative subcomplexes. From a more naïve point of view I would simply state that – because of the underlying data derived from AP-MS experiments – the altProt might be present to a certain extent in the AP-MS but that these two proteins do not interact with each other but, instead, are heterogenous preys (that differ e.g. by 1 AA only) of the bait, so that these comprise a mixture of refPreys and altPreys. In line with this, the networks presented in Figure 3 could also be drawn completely differently. For instance, Figure 3Ai: The hub could be Bend4 but it could also be IP_117582, because in AP-MS the bait protein is sequenced as well and revealed also sequences from IP_117582 upon re-evaluation of the MS data by OpenProt. In my opinion a single circle with two colours (e.g. blue and red) could reflect the findings much more accurately than generating a novel network of interactions based upon the assumption that altProts interact with refProts (which has to be proven anyway in cell-biological or biochemical experiments). In this respect, it would be also beneficial if it were clearly stated in Figure 3 which protein was used as bait. For instance, in Figure 3Aix, is ZNF703 the bait? IP_163248 is the corresponding altProt, but does it form links to hubs/bait proteins? Are these hubs/baits and the preys then different? I consider that this should be better illustrated.

Another example that is not quite clear to me is Figure 3Ci: GML as bait pulls down GAPDH44/IP_761275, which contains only the NAD binding domain of GAPDH? Could this be distinguished by re-evaluation of AP-MS data with OpenProt, i.e.by altPeps when the NAD binding domain are the same (please see also my first point above).

***Response*** *- There are several points here which we address below. We assume this issue primarily relates to pseudogenes-derived protein / parental protein pairs since both proteins may share high similarity levels and peptides specific for the pseudogenes-derived protein may differ from the corresponding parental peptide by one amino acid only. Indeed, altProts encoded in protein-coding genes with an annotated refProt have an amino acid sequence completely different from the refProt*.

**Major comment 8a** - From a more naïve point of view I would simply state that – because of the underlying data derived from AP-MS experiments – the altProt might be present to a certain extent in the AP-MS but that these two proteins do not interact with each other but, instead, are heterogenous preys (that differ e.g. by 1 AA only) of the bait, so that these comprise a mixture of refPreys and altPreys.

***Response*** *- The BioPlex AP-MS experiments were performed with refBaits (reference proteins only were used as baits); hence, the refBait is always identified in the AP-MS because over-expressed. If a prey protein, whether a refPrey or an altPrey is identified in an AP-MS, it typically means that the prey interacts with the refBait. In the case of a parental protein (e.g. the refBait is GAPDH), the identification of the pseudogene-derived protein (e.g. GAPDHP44/IP_761275) in the AP-MS indicates that GAPDHP44/IP_761275 interacts with GAPDH. Obviously, as mentioned by the reviewer, it is possible that heterocomplexes (homomeric & heteromeric) coexist: refBait/refPrey complexes (e.g. GAPDH/GAPDH) and refBait/altPrey complexes (e.g. GAPDH/GAPDHP44-IP_761275). The AP-MS as it was performed would not allow to demonstrate the presence of both types of complexes. However, the conclusion that the altPrey (GAPDHP44/IP_761275) interacts with the refBait (GAPDH), and thus that heterocomplexes with both GAPDH and GAPDHP44/IP_761275 subunits are present in the biological sample remains accurate. The presence of homomeric complexes is possible but remains to be demonstrated using a different experimental strategy, which is out of scope for the current manuscript based on the re-analysis of published AP-MS data*.

**Major comment 8b** - In line with this, the networks presented in Figure 3 could also be drawn completely differently. For instance, Figure 3Ai: The hub could be Bend4 but it could also be IP_117582, because in AP-MS the bait protein is sequenced as well and revealed also sequences from IP_117582 upon re-evaluation of the MS data by OpenProt.

***Response*** *- IP_117582 is an altProt encoded in the BEND4 gene. The altORF is located in the 5’UTR of the annotated mRNAs, upstream of the annotated coding sequence for the Bend4 protein. The refProt and the altProt have 2 completely different amino acid sequences. Hence, our data strongly suggest that IP_117582 is a novel interactor in the Bend4 interactome. In addition, no spectra matching any peptide of the Bend4 protein was found in the whole dataset, only the altProt IP_117582 encoded by BEND4 was found, and the Bend4 protein was not used as bait in BioPlex. Overall, because Bend4 and IP_117582 are two completely different proteins, the possibility that both heteromeric complexes (Bend4/IP_117582) and homomeric complexes (Bend4/Bend4) exist is excessively speculative*.

**Major comment 8c** - In my opinion a single circle with two colours (e.g. blue and red) could reflect the findings much more accurately than generating a novel network of interactions based upon the assumption that altProts interact with refProts (which has to be proven anyway in cell-biological or biochemical experiments).

***Response*** *- In order to acknowledge the possibility that in the case of parental protein / pseudogene-derived protein pairs, both heteromeric (parental protein / pseudogene-derived protein heteromers) and homomeric (parental protein homomers) complexes could exist, we’ve added the following sentence in the legend of Figure 3A: “**Note that in the case of a pseudogene-derived protein in the interactome of its parental protein (iv, v, vii, viii), it is possible that in addition to heteromeric complexes containing both the parental refProt (bait) and pseudogene-derived altProt (prey), homomeric complexes containing at least two subunits of the parental protein (bait and prey) also exist.**”*

**Major comment 8d** - In this respect, it would be also beneficial if it were clearly stated in Figure 3 which protein was used as bait. For instance, in Figure 3Aix, is ZNF703 the bait? IP_163248 is the corresponding altProt, but does it form links to hubs/bait proteins? Are these hubs/baits and the preys then different? I consider that this should be better illustrated.

***Response*** *- Baits are identified with a dark blue colour as indicated in the figure but are not identified by their accession numbers for clarity purposes because some altProts were identified in the interactome of dozens refProts. The subnetworks shown in Figure 3 are meant to present the variety of topology surrounding altProt and we deemed labeling of all baits out of scope. We are preparing a web application that will provide access to all the details of protein clusters and altProt second neighbourhoods:* http://openprot.org/ppi*. This webpage is under construction and will be up and running within the next 4 weeks*.

**Major comment 8e** - Another example that is not quite clear to me is Figure 3Ci: GML as bait pulls down GAPDH44/IP_761275, which contains only the NAD binding domain of GAPDH? Could this be distinguished by re-evaluation of AP-MS data with OpenProt, i.e.by altPeps when the NAD binding domain are the same (please see also my first point above).

***Response*** *- In the GML pull down, peptides identifying the protein IP_761275 encoded by GAPDHP44 map uniquely to that protein and are not shared with the canonical protein encoded by GAPDH. We will include these spectra in a revised manuscript*.

**Major comment 9** - The illustration of the clustering is also not clear to me, mainly for the reasons stated above: For instance, if the authors state that the altProt IP_293201 from gene RNF215 is a novel interactor of the RNA exosomes. Does this mean that only IP_293201 was identified upon re-evaluation of the Bioplex 2.0 data, or both, namely RNF215 and IP_293201? Also, I wonder where in the cluster #15 are the U2 snRNP B’’ protein (SNRPB2) and the U1 snRNP A (SNRPA) are. Both these interact with SNRPA1 for sure.

***Response*** *- Only IP_293201 encoded by the gene RNF215 was identified and not the canonical RNF215 protein*.

*The markov clustering algorithm only takes into account node connectivity when subdividing the network into clusters. While overall cluster composition correlates with known complexes in general, the extracted clusters are not perfect representations of currently known complexes. In BioPlex 2.0 SNPRA does not appear in the interactions of SNRPA1, but it doesn’t mean that they are not interactors, only that they were not detected in those conditions*.

**Major comment 10** - The validation experiments do not entirely convince me. These result from expression of exogenous gene constructs. Here, at least the documentation of the expression level of tagged proteins compared among each other and with the expression of the endogenous proteins is missing.

***Response*** *- Here, we have identified several altProts and we have selected a few of them for further validation. While it is certainly doable to produce custom antibodies to detect the endogenous altProt when focusing on a single specific altProt as we (PMID: 33497625, 33226175, 30181344) and others (PMID: 33535099, 32958672, 27918561) have done before, such an approach is not possible for several novel proteins. When investigating several novel proteins for which no antibodies are commercially available yet, a strategy using protein tags is generally used in the literature to validate the localization and some interactions. Hence, we consider that comparing the expression level of tagged proteins with the expression of the endogenous proteins is beyond the scope of the current manuscript. Obviously, we will raise custom antibodies for the altProts we will focus on in the following manuscripts*.

**Major comment 10a** - In the case of FADD/IP_198808 I admit that I do not see any cytoplasmic localization of FADD in filaments compared with the DAPI staining of the DNA. The shapes of DAPI and FADD staining look similar. The staining from IP_198808 indeed looks different, but this might alternatively be due to the GFP tag and hence different localization. Here, more compelling fluorescence experiments are necessary.

***Response*** *- In the original manuscript showing a typical cell co-transfected with Flag-FADD and IP_198808-GFP, the intensity of the FADD filaments in the cytoplasm was indeed difficult to see compared to the nuclear filaments. We already have and will provide in a revised manuscript new pictures where cytoplasmic FADD filaments are more easily visible*.

**Major comment 10b** - I also recommend a reverse IP, i.e. with anti GFP but also with anti-Flag and vice versa, for all Co-APs/IPs - I recommend that the authors deposit the full WBs.

***Response*** *- We have already performed these experiments and will add them in a revised manuscript*.

**Major comment 10c** - Regarding prohibitin, this is known to oligomerise, so the observed interaction between the refProt and altProt is expected upon expression, because of its interaction within the coil-coiled domains. I suspect that the difference in the AP-MS might have derived from the different tags. I also miss the controls, i.e. Flap, GFP tag alone described in the MM section.

**Response** - AltProt IP_762813 is coded by a prohibitin pseudogene (PHBP19) and the protein is predicted to contain several signatures specific to prohibitin proteins, as indicated in OpenProt: https://www.openprot.org/p/altorfDbView/79/43652080/762813/IP_762813/2/predictedDomainInfo

*As noted by the reviewer, an interaction between the refProt and the altProt was indeed expected but had to be experimentally validated. Our result confirmed such interaction. We consider this result to be very significant since it indicates that a gene annotated as a pseudogene actually encodes a protein that interacts with a complex formed by parental proteins*.

*We do not understand the comment “I suspect that the difference in the AP-MS might have derived from the different tags”. The blot clearly shows a specific interaction between the refProt and the altProt*.

*As for the control for that experiment, the result shows that PHB1-GFP does not bind non-specifically to Flag beads. Furthermore, each AP-MS (Flag pull-down of IP_762813 and GFP pull-down of PHB) were scored and filtered using beads-specific cRAPome using the SAINT algorithm (see Material and Methods, section “Highly confident interacting proteins (HCIPs) scoring of in-house affinity purifications”). Thus, the different tags are highly unlikely to influence the high confidence set of interactors reported for each bait*.

**Major comment 10d** - RPL8 is a ribosomal protein – so it seems remarkable that location of the expressed and tagged protein is exclusively in the nucleus. The authors may wish to comment on this.

***Response*** *- The RPLP8-GFP fusion with the human sequence was previously used as one of the 5891 baits for the large-scale analysis of the human interactome. That is why we used that construct to validate the interaction between RPL18 and IP_117582. We also took advantage of that GFP construct to test the co-localization of both proteins*.

*Although we could observe some fluorescence in the cytoplasm, it was very weak compared to the fluorescence in the nucleus. The nuclear localization, more particularly in the nucleolus is similar to what has been previously described by immunofluorescence*.

*However, RPLP18 also localizes in the cytoplasm were mature ribosomes translate proteins. Thus, we have added the following text to briefly comment on that: “**Similar to endogenous RPL18, RPL18-GFP localized to the nucleus, particularly in the nucleolus. However, it did not localize in the cytoplasm similar to endogenous RPLP18. Thus, it is possible that the GFP tag partially prevents RPL18P from accumulating in the cytoplasm. However, this effect of the GFP tag would not explain the co-localization between RPL18-GFP and IP_117582, or the interaction between both proteins”***.

**Major comment 11** - The authors provided a link under which they have deposited their own AP-MS data (Protein interaction AP-MS data for both IP_762813 and PHB1 in HEK293 cells were deposited to the ProteomeXchange Consortium via the PRIDE (Perez-Riverol et al, 2016) partner repository with the dataset identifier PXD022491. However, the other did not provide any information of how to access these data. I recommend to add these information in a revised version so that the referee(s) can access and evaluate these AP-MS data.

***Response*** *- Here are the login details for PXD022491:*

*Username:*

*Password: PaLFvjZh*

**Minor comment** - The numbers of identified altProts listed in the Results section are different from those in the abstract. Also, the description in the text of the result section is confusing – here, I would appreciate more clarity regarding which and how many altProts including their origin genes (RNAs) the authors have identified as being expressed.

***Response*** *- The confusion may have come from the fact that the abstract indicates the number of alternative proteins within distinct communities only, a number that is not mentioned in the Results section. Thus, we have modified one sentence in the abstract: Here we used the proteogenomic resource OpenProt and a combined spectrum- and peptide-centric analysis for the re-analysis of a high throughput human network proteomics dataset thereby revealing the presence of <u>280</u> alternative proteins <u>in the network</u>*.

